# Setdb2 Regulates Inflammatory Trigger-Induced Trained Immunity of Macrophages Through Two Different Epigenetic Mechanisms

**DOI:** 10.1101/2025.03.18.644013

**Authors:** Anant Jaiswal, Laszlo Halasz, David L Williams, Timothy Osborne

**Author notes:** Research & Education Building, 600 5th Street South, St. Petersburg, FL 33701. Senior author (Timothy Osborne). bluesky:@timtocool.bsky.social (Timothy Osborne).

## Abstract

"Trained immunity" of innate immune cells occurs through a sequential two-step process where an initial pathogenic or sterile inflammatory trigger is followed by an amplified response to a later un-related secondary pathogen challenge. The memory effect is mediated at least in part through epigenetic modifications of the chromatin landscape. Here, we investigated the role of the epigenetic modifier Setdb2 in microbial (β-glucan) or sterile trigger (Western-diet-WD/oxidized-LDL-oxLDL)-induced trained immunity of macrophages. Using genetic mouse models and genomic analysis, we uncovered a critical role of Setdb2 in regulating proinflammatory and metabolic pathway reprogramming. We further show that Setdb2 regulates trained immunity through two different complementary mechanisms: one where it positively regulates glycolytic and inflammatory pathway genes via enhancer-promoter looping, and is independent of its enzymatic activity; while the second mechanism is associated with both increased promoter associated H3K9 methylation and repression of interferon response pathway genes. Interestingly, while both mechanisms occur in response to pathogenic training, only the chromatin-looping mechanism operates in response to the sterile inflammatory stimulus. These results reveal a previously unknown bifurcation in the downstream pathways that distinguishes between pathogenic and sterile inflammatory signaling responses associated with the innate immune memory response and may provide potential therapeutic opportunities to target cytokine vs. interferon pathways to limit complications of chronic inflammation.

## INTRODUCTION

The vertebrate innate immune system has evolved a robust mechanism that recognizes prior pathogen exposure and directs a rapid and robust secondary response to a broad spectrum of infectious agents to limit deleterious effects on organismal health and survival. This learned response has been described as “trained immunity” or “innate immune memory” and provides protection from a broad range of potential future pathogens.^1^ Specific molecular features associated with disease-causing microbes have been identified that initiate the process, and they range from microbial surface carbohydrates such as β-glucan to engineered mimics including the bacillus Calmette-Guérin (BCG) vaccine.^2,3^ However, endogenous triggers that arise during “sterile” inflammatory challenges, for instance, Western diet (WD) feeding and some intermediary metabolites, can also induce a trained immune response.^4^ The process occurs at least in part through epigenetic reprogramming of chromatin in hematopoietic stem cells, dendritic cells, monocytes, and macrophages linked to genes involved in modulating inflammatory and supporting metabolic pathways.^5,6^

The capacity to mount a second proinflammatory challenge response after the primary exposure inherent to the innate immune memory system are retained over time and can be inherited following cell division, which suggests they are influenced by metastable epigenetic modifications. Consistent with this model, recent reports have revealed the importance of key histone modifications and their modifier enzymes in the mechanism. For instance, mono- or tri-methylation at lysine four of histone H4 (H3K4me1 and H3K4me3) within enhancers and promoters of affected genes contributes to β-glucan-induced trained immunity,^7, 8, 9^ and the response was inhibited in macrophages lacking *Set7*, an enzyme regulating H3K4me1.^10^ Additionally, treatment of cells with pan-methyltransferase inhibitors also resulted in a decrease in the innate immune memory response.^5^ An additional study highlighted the role of the H3K9me2 (di-methylation at lysine-9 position of histone) modifying enzyme G9a as a regulator of trained immunity.^11^ Despite these specific examples, how specific chromatin-dependent modifications influence the training process and whether there is heterogeneity in the downstream secondary responses that are related to the identity or source of the initiating trigger have not been explored.

SET domain bifurcated 2 (Setdb2) is a member of the SET domain-containing KMT1 subfamily of lysine methyltransferases.^12^ The SET domain binds S-adenosyl methionine (SAM), which is a methyl-donating cofactor for virtually all known lysine methyltransferases. KMT1 family members such as SETDB1, G9a, and SUV3-9H1 utilize SAM to methylate histone H3 on lysine 9 (H3K9).^12,13^ H3K9 methylation is a prominent gene-repressive histone mark, and recent studies have shown Setdb2 is induced early during a proinflammatory response in macrophages and is required for the increase in H3K9me3 that occurs at its target cytokine gene-promoters to decrease their expression and limit runaway inflammation.^14^ Setdb2 was also shown to be involved in regulating H3K9 methylation in diabetic wound-associated macrophages in mice.^15^

In a totally unrelated investigation, we uncovered a different role for Setdb2 in hepatocytes, where we showed that Setdb2 was induced by glucocorticoid receptor (GR) signaling during fasting. We showed that Setdb2 works together with GR to modulate long-range enhancer-promoter interactions in hepatic chromatin to stimulate Insig2 gene expression and repress sterol regulatory element binding protein (SREBP)-dependent lipogenesis during fasting.^16^ Additionally, we found that coactivation of GR target genes by Setdb2 was independent of its putative methyltransferase activity and was associated with a decrease in H3K9 methylation of target gene promoters. In a follow-up study, we also found that Setdb2 was highly expressed in human atherosclerotic lesions, and transplantation of bone marrow from global Setdb2-deficient mice into *Ldlr-/-* mice resulted in increased plaque-associated vascular inflammation with altered immune cell profiles and accelerated atherosclerotic progression. This suggests Setdb2 is involved with chronic inflammation associated with metabolic disorders such as atherosclerosis.^17^

Here, we investigated the role of Setdb2 in response to microbial (β-glucan) or endogenous sterile inflammatory stimuli Western-diet (WD) and oxidized-LDL (oxLDL)) models of the innate immune memory response in macrophages challenged in vivo or in vitro. We compared responses in WT versus two key genetic models: one where Setdb2 is specifically deleted in myeloid cells and another where we knocked in alanine point mutations to alter two key amino acids in the SET domain that are key to coordinate binding to SAM for its methyltransferase activity. The results show that Setdb2 is critical for the induction of trained immunity in macrophages triggered both in vivo and in vitro and in response to pathogenic and sterile inflammatory signals. We combined gene expression analyses with chromatin immunoprecipitation (ChIP) and chromatin conformation capture (3C) analyses to show that Setdb2 plays a central role in regulating proinflammatory and metabolic pathway genes through two different mechanisms. In one, Setdb2 positively regulates glycolytic and inflammatory cytokine pathway genes via changes in enhancer-promoter looping, and this requires Setdb2 but is independent of its enzymatic activity. In the second mechanism, Setdb2 enzyme activity and its SET domain is required to negatively regulate genes involved in inflammatory and interferon pathway signaling, and this is associated with H3K9 methylation changes in target gene promoters. These results reveal a bifurcation of the innate immune memory response into pathways that, on the one hand, activate pro-inflammatory and metabolic responses while inhibiting interferon-associated signaling pathways. They also reveal a divergent memory response initiated by β-glucan vs WD/ox-LDL, which is consistent with different training responses downstream from infectious vs sterile chronic inflammatory triggers.

## RESULTS

### Setdb2 is essential for β-glucan-induced innate immune memory of bone-marrow macrophages

To investigate the role of Setdb2 in innate immune memory, we first established a two-step in vitro culture model of trained immunity in bone marrow (BM)-macrophages exposed to the yeast cell wall carbohydrate β-glucan as a prototype pre-training signal as described by others^10^ and Figure 1A. Briefly, freshly isolated mouse bone marrow cells were incubated with β-glucan or culture medium. After 24 h, cells were washed and cultured with GM-CSF for an additional five days and then treated with culture medium alone or medium supplemented with lipopolysaccharide (LPS) for six hours. The culture supernatant was collected, and cells were harvested for RNA and protein analysis (Figure 1A). Following this protocol, we found that untrained LPS treatment alone induced *Setdb2* mRNA expression. Interestingly, LPS stimulation effect was further amplified in WT cells that were pre-trained with β-glucan in WT (Figure 1B). Setdb2 protein levels followed a similar pattern, and as expected, Setdb2 mRNA and protein expression was negligible in BM-macrophages isolated from mice with a myeloid-specific knockout of Setdb2 (Setdb2mKO) (Figure 1B, C). Thus, the expression of Setdb2 was super-induced by β-glucan training similar to classic trained immunity responses observed in previous reports.^10^

**Figure 1.**
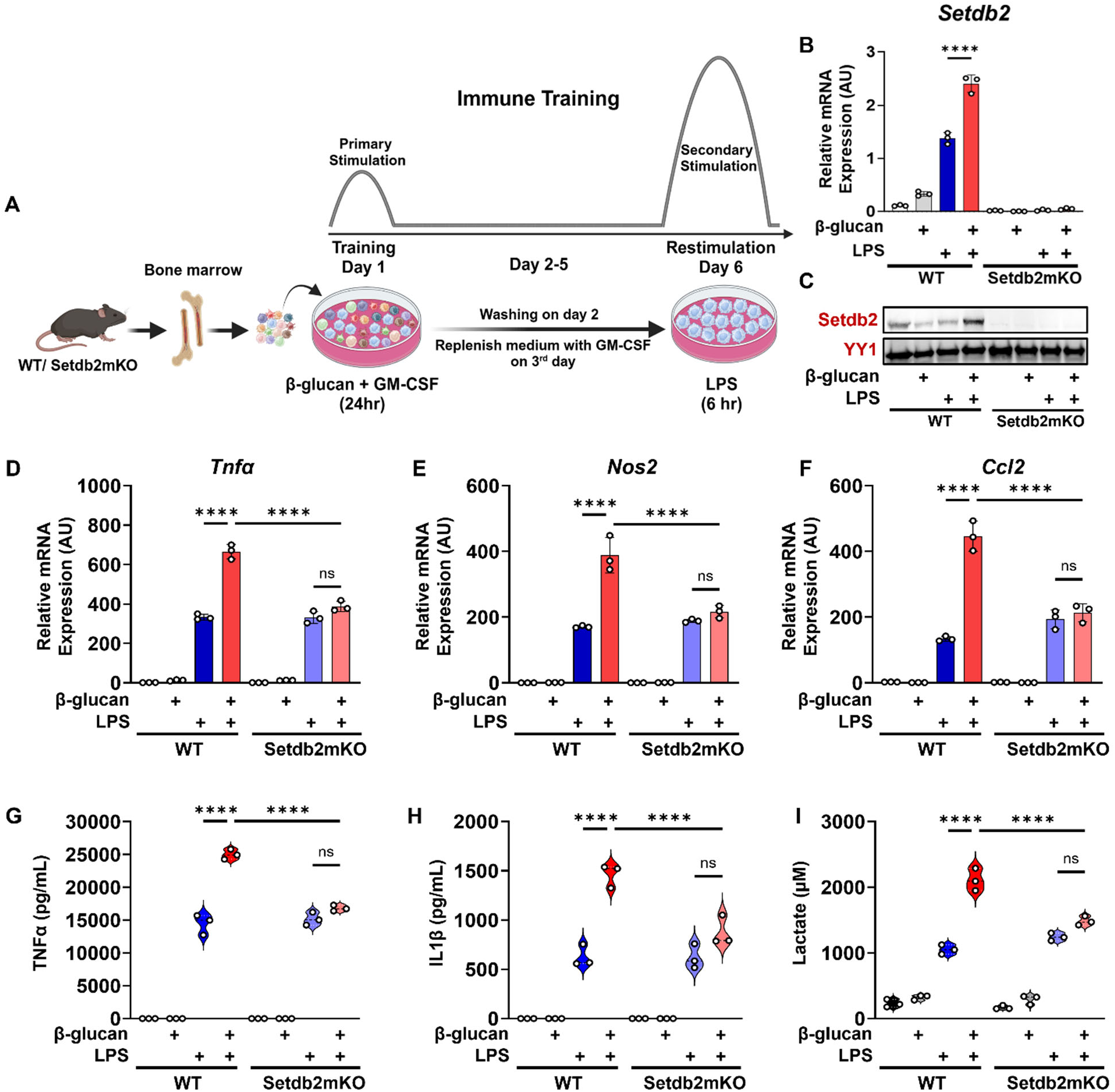
Setdb2 is essential for β-glucan induced innate immune memory of bone-marrow macrophages. (A) Graphical outline showing in vitro β-glucan training model. Bone marrow cells were isolated from WT and Setdb2mKO mice and treated with 5 μg/mL β-glucan (training stimulus) or culture medium for 24 h, cultured for 5 days, and restimulated with 100ng/mL LPS (secondary challenge) or culture medium for 6 h on day 6, cells were harvested for analysis. (B) Setdb2 mRNA expression and (C) Setdb2 protein expression, showing one representative immunoblot from three independent experiments. YY1 serves as a loading control, (D) mRNA expression of Tnfα (E) Nos2, and (F) Ccl2, (G) Pro-inflammatory cytokine levels of TNFα, and (H) IL-1β, and (I) Extracellular lactate levels in untreated control, β-glucan alone, LPS alone, and β-glucan followed by LPS restimulated BM-macrophages from WT (n = 3) and Setdb2mKO mice (n = 3). Data are represented as mean ± SD. ‘n’ represents biological replicates of each strain and treatment type. p-value was calculated using two-way ANOVA with Tukey post-hoc test for multiple comparisons. ∗p < 0.05, ∗∗p < 0.01, and ∗∗∗p < 0.001, ns, not significant change.

To explore the role of Setdb2 in the training mechanism, we measured expression of mRNAs for key proinflammatory markers of the classic innate immune memory response such as nitric oxide synthase (*Nos2*), tumor necrosis factor (*Tnfα*) and monocyte chemoattractant protein 1 (*Ccl2*). These genes were induced by untrained LPS alone treatment and super-induced by β-glucan pre-training in WT similar to Setdb2 expression as expected. Interestingly, the super-induction by β-glucan pre-training was significantly blunted in BM-macrophages isolated from Setdb2mKO mice (Figure 1D-F). Similarly, secretion of Tnfα and IL1β cytokines from the cells was super-induced in WT, but this was also significantly blunted in Setdb2mKO BM-macrophages (Figure 1G-H). During trained immunity of macrophages, cells also shift their metabolism to increase aerobic glycolysis,^5^ so we also measured extracellular lactate levels as a readout of this metabolic shift. This response was also enhanced by pre-training in WT, and it was significantly reduced in Setdb2mKO BM-macrophages (Figure 1I). Together these findings show that Setdb2 plays a critical role in the innate immune memory response of BM-macrophages.

### Setdb2 regulates key metabolic and inflammatory pathways during β-glucan-induced trained immunity in bone-marrow macrophages

To explore the effects of Setdb2 on the global transcriptional program during the trained immune response, we performed whole-genome RNA sequencing of β-glucan-pre-trained WT vs. Setdb2mKO BM-macrophages. Principal component analysis revealed that control cells from WT and Setdb2mKO mice clustered together, which is consistent with Setdb2 not having a dominant role in unstimulated cells. However, the PCA patterns were shifted for all treatment groups, and there were clear differing patterns for cells from WT vs. Setdb2mKO, consistent with a key role of Setdb2 in the training process (Figure 2A). Over 2000 genes were differentially regulated comparing β-glucan pre-trained versus untrained LPS-treated BM-macrophages, as shown by the volcano plots in Figure 2B and more detailed in supplemental table S2. More than 1000 of these were differentially expressed in β-glucan pre-trained cells followed by challenged with LPS from WT vs Setdb2mKO mice (Figure 2C).

**Figure 2.**
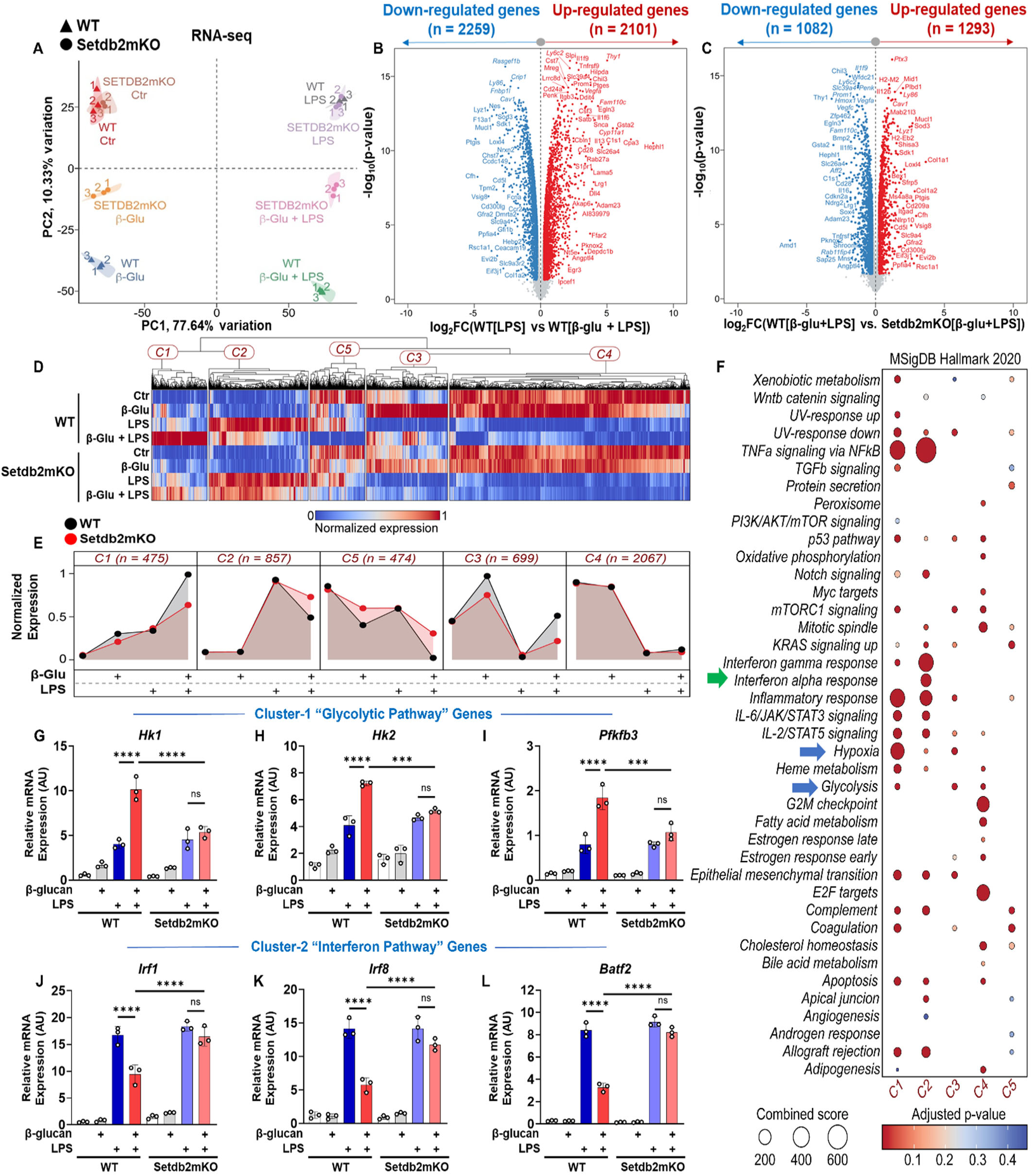
Setdb2 regulates key metabolic and inflammatory pathways during β-glucan-induced trained immunity in bone-marrow macrophages. Bone marrow cells were isolated from WT and Setdb2mKO mice and treated with 5 μg/mL β-glucan (training stimulus) or culture medium for 24 h, cultured for 5 days, and restimulated with 100ng/mL LPS (secondary challenge) or culture medium for 6 h on day 6, cells were harvested and analyzed by RNA-sequencing. (A) Principal component analysis (PCA) showing distinct transcriptional profiles across treatment conditions and genotypes (n = 3). (B) Volcano plot visualizing differential gene signatures in cells trained with β-glucan followed by LPS restimulation vs LPS alone treated BM-macrophages from WT (n = 3). (C) Volcano plot visualizing differential gene signatures in cells trained with β-glucan followed by LPS restimulation for WT vs Setdb2mKO BM-macrophages (n = 3). (D) Heatmap, and (E) Area plot showing k-means (k = 5) and hierarchical clustering (n = 3). ("See also Figure S1") (F) Enrichment analysis using MSigDB to identify significantly enriched biological processes associated with the DEGs across experimental groups (n = 3). (G) mRNA expression of Hk1 (H) Hk2, (I) Pfkfb3, (J) Irf1, (K) Irf8, and (L) Batf2 in untreated control, β-glucan alone, LPS alone, and β-glucan followed by LPS restimulated BM-macrophages from WT (n = 3) and Setdb2mKO mice (n = 3). Statistical analysis for RNA-seq data was performed using DESeq2 with its default settings. Data are represented as mean ± SD. ‘n’ represents biological replicates of each strain and treatment type. p-value was calculated using two-way ANOVA with Tukey post-hoc test for multiple comparisons. ∗p < 0.05, ∗∗p < 0.01, and ∗∗∗p < 0.001, ns, not significant change.

Unsupervised k-means clustering revealed a broad pattern of transcriptional changes that were distributed into 5 unique clusters (k=5) visualized by the heatmaps in Figure 2D and Figure S1A-E. The cluster effects were pooled together to generate a cluster-wide response using a line plot format that emphasizes the dynamic nature of the transcriptional profiles for the five clusters. Importantly, the expression pattern for cluster-1 (475 genes) and cluster-2 (857 genes) displayed interesting transcriptional profiles with respect to Setdb2 loss and β-glucan pre-training. Genes associated with cluster-1 were induced by LPS treatment alone and super-induced by β-glucan pre-training followed by LPS stimulation in WT, while this super-induction was significantly blunted in Setdb2-deficient BM-macrophages. In contrast, cluster-2 associated genes were induced by LPS, but the LPS-mediated induction was inhibited by β-glucan pre-training in WT. However, this inhibition by β-glucan pre-training was lost in the Setdb2mKO BM-macrophages.

To explore the biological processes and molecular pathways associated with the transcriptional profiles, we performed a gene enrichment analysis using Molecular Signatures Database (MSigDB) on the genes contained in all five clusters (Figure 2F). This analysis showed that the majority of genes from cluster-1 and cluster-2 were associated with proinflammatory and metabolic pathways in comparison with other clusters. Cluster-1 genes were specifically enriched in proinflammatory and metabolic pathways including hypoxia and glycolysis, while cluster-2 genes were preferentially associated with interferon alpha and gamma pathways. Some proinflammatory pathways, such as JAK/STAT, as well as TNFα signaling via *NFκB,* were equally distributed within both cluster-1 and 2 enriched genes. Genes in the other clusters were associated with other processes, and because their overall response to treatment and genotype was more complex, we focused on clusters 1 and 2.

To validate our RNA-sequencing data, we performed qPCR analysis to measure key genes in clusters 1 and 2. Consistent with the RNA-seq data, we found that cluster-1-associated glycolytic genes, *i.e.*, *Hk1, Hk2, and Pfkfb3,* were significantly induced by LPS and super-induced by β-glucan pre-training in WT, similar to *Tnfα, Nos2, and Ccl2*, and this response was blunted in Setdb2mKO BM-macrophages (Figure 2G-I). Similarly, consistent with the RNA-seq results, cluster-2-associated interferon pathway genes, *i.e.*, *Irf1, Irf8, and Batf2,* were induced by LPS but this was inhibited by β-glucan pre-training in WT, but the inhibition was also lost in Setdb2mKO BM macrophages (Figure 2J-L). Taken together, these results confirm that Setdb2 is required for both β-glucan training-dependent activation of cluster-1 genes and β-glucan training-dependent suppression of cluster-2 genes.

### Setdb2 regulates in vivo β-glucan induced trained immunity in peritoneal macrophages

To validate our *in vitro* findings further and to analyze the effect of β-glucan pre-training *in vivo*, β-glucan was injected into the peritoneal cavity of WT and Setdb2mKO mice. After 6 days we isolated the peritoneal macrophages and treated them acutely with LPS as a secondary stimulus (diagrammed in Figure 3A). Similar to the β-glucan in-vitro challenge, we found that expression of proinflammatory genes in cluster-1 including *Tnfα, Nos2,* and *Ccl2* and secretion of Tnfα and IL-1β cytokines were super-induced by β-glucan training in vivo, and these responses were blunted in Setdb2mKO peritoneal macrophages (Figure 3B-F). There was a similar effect on lactate production and the expression of cluster-1 associated glycolytic genes, i.e., *Hk1, Hk2,* and *Pfkfb3* (Figure 3G-J). In contrast, but similar to our in vitro training studies, the LPS induction of cluster-2 interferon pathway genes (i.e., *Irf1, Irf8, Batf2*) was blunted by β-glucan pre-training in WT, and this was not observed in Setdb2mKO (Figure 3K-M). Overall, our data provide both in vitro and in vivo evidence that Setdb2 regulates β-glucan training-mediated induction and repression of cluster 1 and 2 genes, respectively.

**Figure 3.**
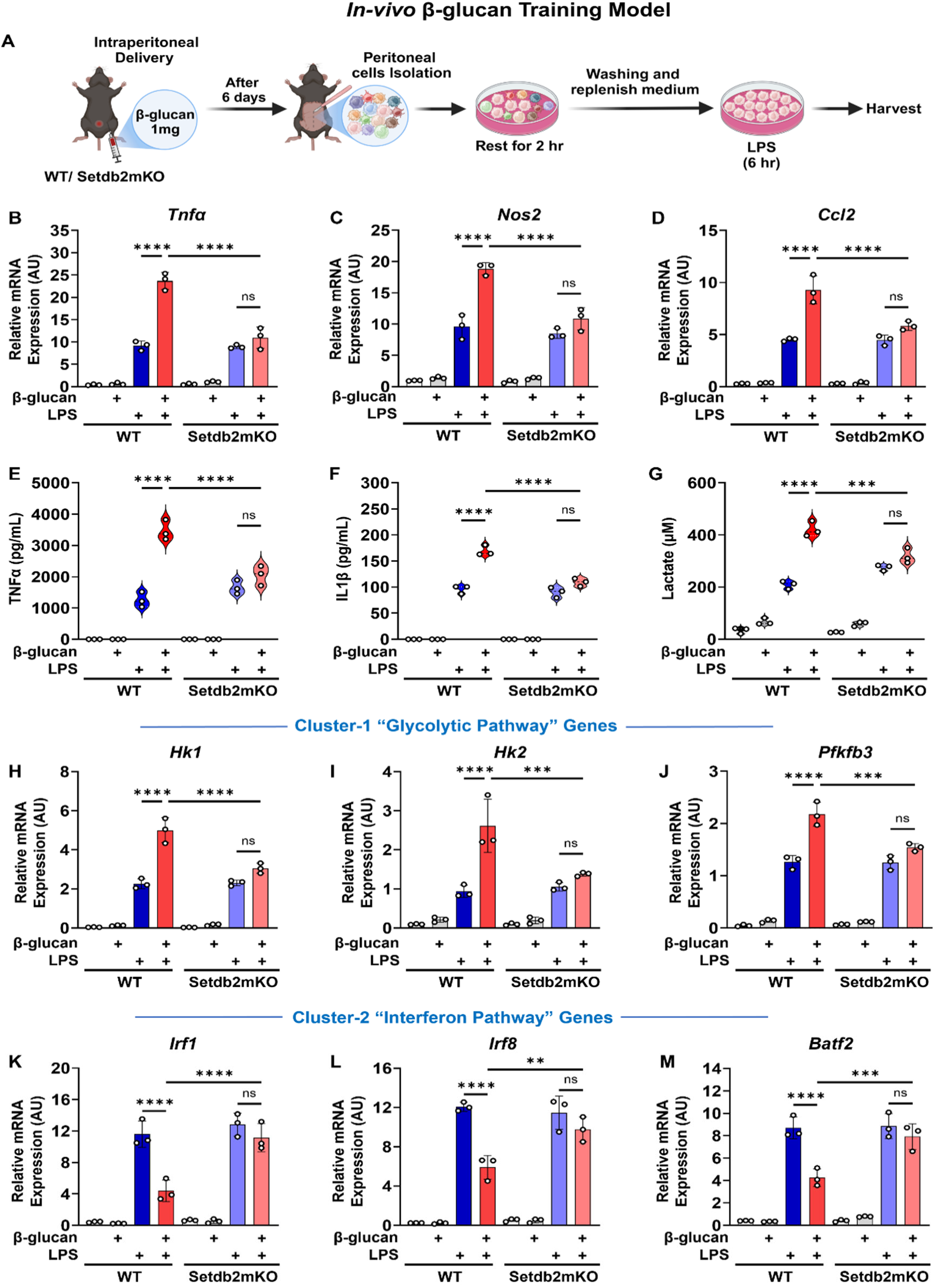
Setdb2 regulates in-vivo β-glucan induced trained immunity in peritoneal macrophages. (A) Graphical outline showing in vivo β-glucan training model. WT or Setdb2mKO mice were peritoneally injected with 1mg β-glucan (training stimulus) or PBS alone and after 5 days peritoneal macrophages were isolated and restimulated with 100ng/mL LPS (secondary challenge) or culture medium for 6 h. Cells were harvested for analysis. (B) mRNA expression of Tnfα, (C) Nos2, and (D) Ccl2, (E) Pro-inflammatory cytokine levels of TNFα, and (F) IL-1β, (G) Extracellular lactate levels, (H) mRNA expression of Hk1 (I) Hk2, (J) Pfkfb3, (K) Irf1 (L) Irf8, and (M) Batf2 in untreated control, β-glucan alone, LPS alone, and β-glucan followed by LPS restimulated peritoneal macrophages from WT (n = 3) or Setdb2mKO mice (n = 3). Data are represented as mean ± SD. ‘n’ represents biological replicates of each strain and treatment type. p-value was calculated using two-way ANOVA with Tukey post-hoc test for multiple comparisons. ∗p < 0.05, ∗∗p < 0.01, and ∗∗∗p < 0.001, ns, not significant change.

### β-glucan-dependent trained immunity in BM-macrophages from Setdb2 knock-in mice

Setdb2 is a member of the KMT1 family of lysine methyltransferases, which includes several well-characterized enzymes that methylate lysine 9 on histone 3 to repress transcription.^13^ Consistent with this notion, recent studies have suggested that Setdb2 is linked with gene repression, where it is associated with an increase in H3K9me3 levels in the promoters of genes that are proposed to be suppressed by Setdb2 during proinflammatory stimulation in macrophages.^14,15^ However, the differential effect of Setdb2 on the response patterns for cluster-1 vs. cluster-2 genes in the current study suggests Setdb2 may regulate cluster-1 positively while inhibiting expression of cluster-2 genes in response to β-glucan training. To explore whether Setdb2’s methyltransferase activity might be involved in differential regulation of cluster-1 vs. cluster-2 pathway genes, we developed a knock-in mouse line (Setdb2Kin), which has germline mutations that change two critical amino acids of the conserved SET domain to alanine. Based on the structure of the SET domain bound to SAM,^12^ these mutations will prevent binding of this required cofactor and eliminate any direct methyltransferase activity (Figure 4A). We found the knock-in mice were viable, fertile, and showed no overt signs of lethality when grown under standard laboratory conditions. Sanger sequencing further confirmed the two-point mutations were introduced into the SET domain of the Setdb2 gene in Setdb2Kin mice (Figure S2A-C). Next, we compared BM-macrophages from the Setdb2Kin to WT mice in the β-glucan training protocol. Interestingly, unlike in Setdb2mKO, we found that the LPS-dependent induction of mRNAs encoding cluster-1 proinflammatory genes in Setdb2Kin such as *Tnfα, Nos2,* and *Ccl2* were still super induced by β-glucan pre-training similar to WT (Figure 4B-D). Similar results were observed for the secretion of TNFα, IL-1β, and lactate levels (Figure 4E-G). Cluster-1 associated glycolytic pathway genes *Hk1, Hk2*, and *Pfkfb3* were also super-induced by β-glucan pre-training in the Setdb2Kin similar to WT (Figure 4H-J). In contrast, the β-glucan dependent repression of the LPS-induction of cluster-2-associated interferon pathway genes, *Irf1, Irf8*, and *Batf2,* was blocked in Setdb2Kin BM-macrophages (Figure 4K-M), which was similar to results for Setdb2mKO. Taken together, these results suggest that Setdb2 may have different roles in regulating cluster-1 and cluster-2 associated pathways.

**Figure 4.**
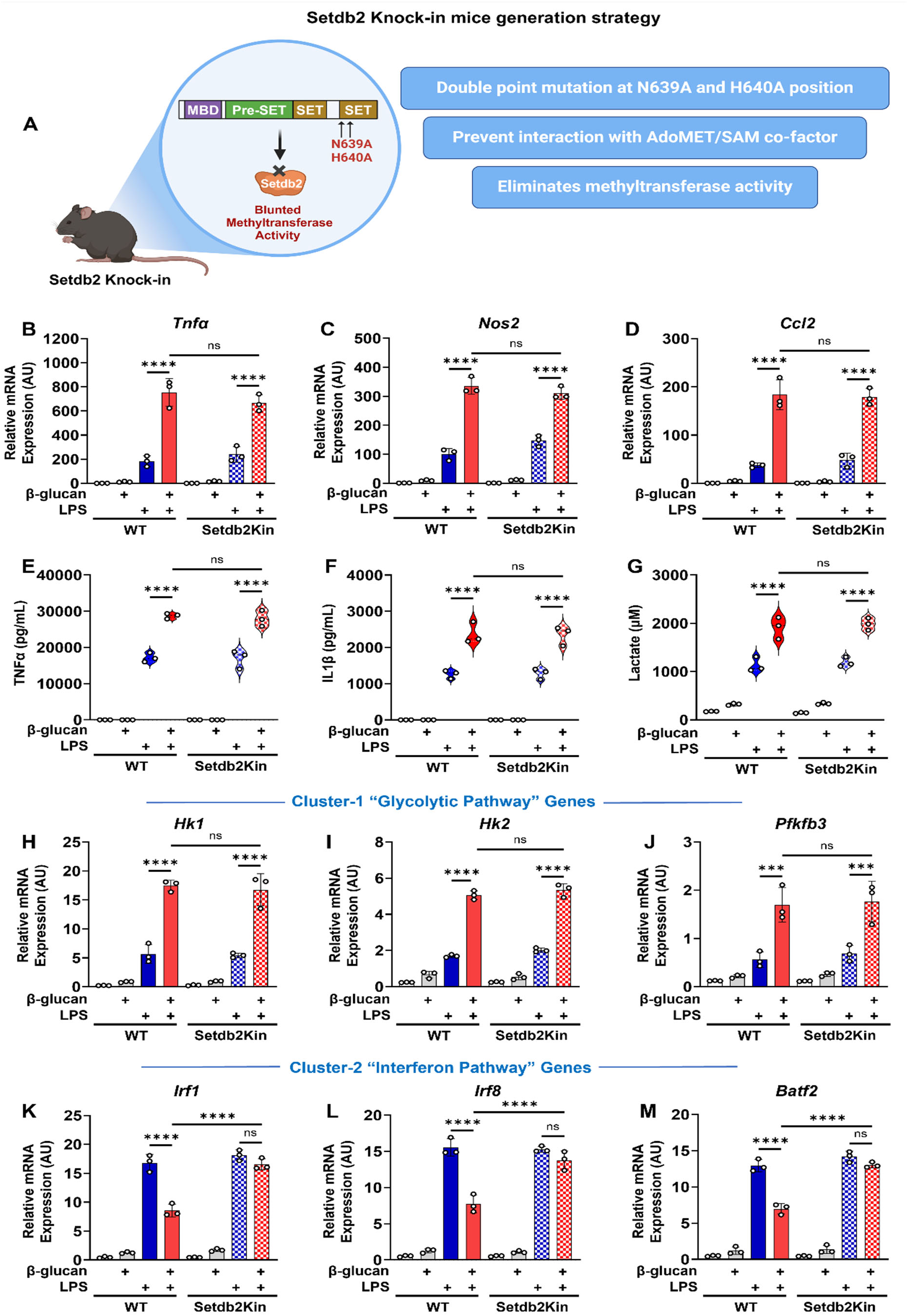
β-glucan-dependent trained immunity in BM-macrophages from WT vs. Setdb2 knock-in mice. (A) Schematic diagram showing Setdb2 knock-in strategy ("See also Figure S2"). Bone marrow cells from WT and Setdb2Kin mice were treated with 5 μg/mL β-glucan (training stimulus) or culture medium for 24 h, cultured for 5 days, and restimulated with 100ng/mL LPS (secondary challenge) or culture medium for 6 h on day 6. Cells were harvested for analysis. (B) mRNA expression of Tnfα, (C) Nos2, and (D) Ccl2, (E) Pro-inflammatory cytokines TNFα, and (F) IL-1β levels, (G) Extracellular lactate levels, (H) mRNA expression of Hk1 (I) Hk2, (J) Pfkfb3, (K) Irf1 (L) Irf8, and (M) Batf2 in untreated control, β-glucan alone, LPS alone, and β-glucan followed by LPS restimulated BM-macrophages from WT (n = 3) and Setdb2Kin mice (n = 3). Data are represented as mean ± SD. ‘n’ represents biological replicates of each strain and treatment type. p-value was calculated using two-way ANOVA with Tukey post-hoc test for multiple comparisons. ∗p < 0.05, ∗∗p < 0.01, and ∗∗∗p < 0.001, ns, not significant change.

### Setdb2 regulates innate immune memory via two distinct mechanisms: activation through chromatin looping and suppression through its methyltransferase activity

Our lab previously reported that Setdb2 is induced during fasting in the liver, where it functions together with the glucocorticoid receptor (GR) to activate a subset of GR target genes involved in the metabolic adaptation to fasting. In this role, Setdb2 was associated with long-range enhancer-promoter interactions and was independent of its enzymatic activity.^16^ Thus, based on these prior findings and considering the results presented above for Setdb2mKO vs. Setdb2Kin macrophages, we hypothesized that Setdb2 may regulate cluster-1 gene activation through enhancer-promoter looping while repressing cluster-2 genes through its methyltransferase activity.

To test our hypothesis, we first performed chromosome conformation capture (3C-qPCR) to analyze long-range enhancer-promoter interactions adjacent to cluster-1 genes. We targeted the promoter and enhancers for *Tnfα, Ccl2,* and *Hk2,* all cluster-1 genes, in chromatin isolated from BM-macrophages subjected to the in vitro β-glucan training protocol. Our 3C-qPCR data demonstrated a β-glucan pre-training dependent increase in the association between the +8 kb and +18 kb enhancer regions for *Tnfα*, a +23 kb for *Ccl2*, and a +48 kb upstream enhancer region for *Hk2*, with the corresponding transcription start sites (TSS) for each gene in chromatin from WT vs Setdb2mKO BM-macrophages. The data show significant enhancer-promoter interactions that were more robust in chromatin isolated from LPS-treated BM-macrophages that were pre-trained with β-glucan compared to chromatin from untrained samples treated only with LPS. Importantly, this response was significantly lower in chromatin from Setdb2mKO mice BM-macrophages (Figure 5A-C). These results suggest that Setdb2 acts to facilitate enhancer-promoter interactions that influence cluster-1 pathway genes during the β-glucan training response.

**Figure 5.**
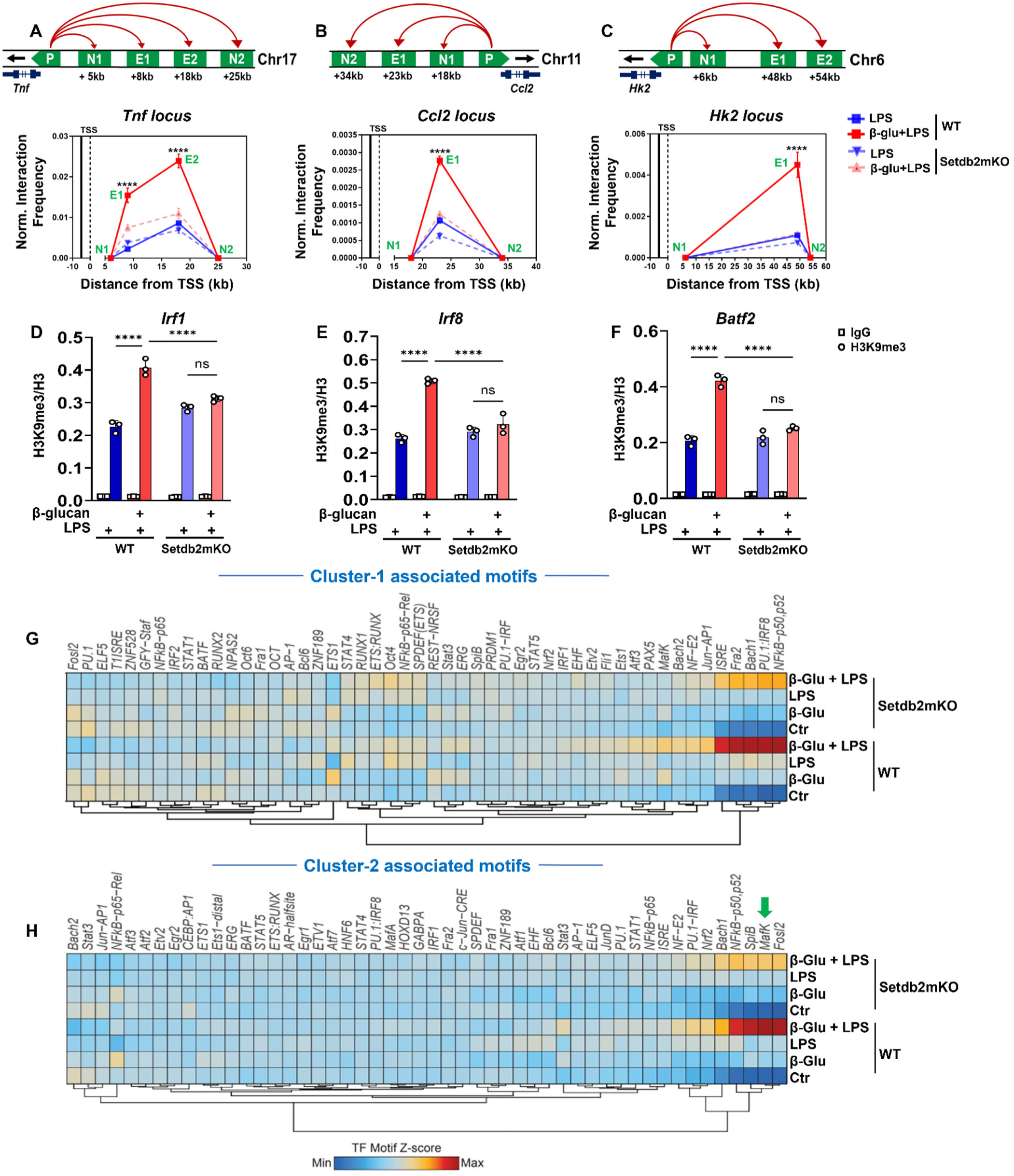
Setdb2 regulates innate immune memory via two distinct mechanisms: activation through chromatin looping and suppression through its methyltransferase activity. Bone marrow cells were isolated from WT and Setdb2mKO mice and treated with 5 μg/mL β-glucan (training stimulus) or culture medium for 24 h, cultured for 5 days, and restimulated with 100ng/mL LPS (secondary challenge) or culture medium for 6 h on day 6. Chromatin was harvested and analyzed by 3C-qPCR, ChIP-qPCR and ATACseq analysis. (A) Diagram representing the chromosomal loci at respective genes, P=promoter region, N= negative bait, E= enhancer bait for potential long-range enhancer-promoter interaction through 3C-qPCR analysis at Tnfα, (B) Ccl2, or (C) Hk2 locus, enhancer bait regions for 3C-qPCR spans the promoter regions from TSS to +8 kb and +18 kb for Tnfα, +23kb for Ccl2, and +48kb regions for Hk2 loci. (D) H3K9me3 levels (ChIP-qPCR) at Irf1 (E) Irf8, and (F) Batf2 promoter in chromatin of LPS alone, or β-glucan followed by LPS restimulated BM-macrophages from WT (n = 3) or Setdb2mKO mice (n = 3). ("See also Figure S3") (G) ChromVAR z-scores heatmap representing significantly enriched HOMER motifs for differentially opened chromatin regions in cluster 1 and (H) cluster-2 gene loci from ATAC-seq data performed with chromatin from untreated control, β-glucan alone, LPS alone, and β-glucan followed by LPS restimulated BM-macrophages from WT (n = 3) and Setdb2mKO mice (n = 3). ("See also Figure S4") For ATAC-seq data, differential peak accessibility was analyzed using DiffBind with EdgeR, employing default parameters. Data are represented as mean ± SD. ‘n’ represents biological replicates of each strain and treatment type. p-value was calculated using two-way ANOVA with Tukey post-hoc test for multiple comparisons. ∗p < 0.05, ∗∗p < 0.01, and ∗∗∗p < 0.001, ns, not significant change.

Next, to determine whether Setdb2 suppression of cluster-2 genes might be mediated through changes in promoter-associated H3K9 methylation, we performed ChIP-qPCR to measure H3K9 methylation levels at the promoters of cluster-2-specific interferon pathway genes. H3K9 tri-methylation levels were significantly increased by β-glucan pre-training at the promoters of *Irf1, Irf8,* and *Batf2* genes in WT mice, as compared to untrained samples treated only with LPS (Figure 5D-F). Interestingly, H3K9 promoter methylation levels were also significantly decreased in Setdb2mKO BM-macrophages. We also evaluated H3K9 mono (Figure S3A-C) and di-methylation (Figure S3D-F) levels and these chromatin marks also followed the same pattern. In contrast, there were no major changes observed with total histone H3 levels at these promoter sites (Figure S3G-I). Overall, these data suggest that cluster-2 pathway gene repression by β-glucan pre-training was associated with elevated levels of promoter-specific H3K9 methylation, which is consistent with a role for Setdb2’s catalytic activity being required for the reduced levels of gene expression. (Figures 5D-F)

To interrogate the chromatin accessibility landscape that is influenced by β-glucan training, we performed Assay for Transposase-Accessible Chromatin sequencing (ATAC-seq) and analyzed the peak data using ChromVar to gain insights into the potential motifs that are associated with differential peaks across the different treatment groups. This analysis showed significant differences in peak distribution comparing β-glucan-trained LPS-challenged WT vs. Setdb2mKO BM-macrophages. A detailed analysis of the motifs associated with variation between the two genotypes showed (Figure S4 B, C) that *NFκB* motif was highly enriched within open chromatin regions associated with both cluster 1 and 2 pathway genes in WT chromatin and the signal was significantly lower in the Setdb2mKO samples (Figure S4A and Figure 5 G-H). This is consistent with our RNA seq data, where *NFκB* signaling genes are present in both clusters. The comparison also showed that the repressor *MafK* motif was enriched within the cluster-2 specific peaks (Figure 5I) which is interesting because MafK has been shown to be recruited along with Setdb1 and the SAM-synthesizing MatIIa/MatIIb dimer to repress target gene expression.^18^

### Setdb2 regulates ‘western-diet-induced’ innate immune training of bone-marrow monocytes

In a previous study, we reported that an adoptive transfer of bone marrow from mice with a global deletion of Setdb2 into irradiated *Ldlr*-knockout mice resulted in increased vascular inflammation and elevated progression of atherosclerosis when the mice were fed a western diet (WD) relative to mice that were given bone marrow from WT mice.^17^ Recent literature also indicates that initial exposure of innate immune cells to endogenous atherogenic stimuli during WD feeding in mice results in long-lasting hyperresponsive immune cells in the bone marrow. This “training” effect in bone marrow cells leads to functional reprogramming, which eventually worsens atherosclerosis.^4^

To determine whether Setdb2 might also be involved in the early reprogramming of bone marrow cell during WD-induced immune training, we fed WT, Setdb2mKO and Setdb2Kin mice with a WD for 20 weeks. This resulted in similar increases in body weight, liver weight, and serum cholesterol levels in all three genotypes (Figure S5A-F). Next, to evaluate whether the loss of Setdb2 affected the pattern of innate immune training in WD-fed mice, we isolated CD11b+ monocytes from the bone marrow of control and WD-fed WT and Setdb2mKO mice and challenged them acutely with LPS (Figure 6A). qPCR analysis for cluster-1 associated pathway genes showed that the WD feeding resulted in a significantly more robust response to LPS challenge compared to control diet-fed mice, consistent with an in vivo training effect in WD-fed WT mice (Figure 6B-D). However, similar to the results for β-glucan training, this enhanced response was not observed in Setdb2mKO monocytes. The expression of key cluster 1-specific glycolytic genes and the secretion of cytokines TNFα and IL1β and lactate were all similarly affected (Figure 6E-J). Consistent with the β-glucan training results, the WD-dependent super-induction following LPS stimulation was dramatically blunted in monocytes from Setdb2mKO mice. Interestingly, in contrast to our observations where β-glucan pre-training resulted in suppression of the LPS activation of cluster-2 specific interferon pathway genes, WD feeding resulted in significant induction of these genes by LPS in both WT and Setdb2mKO samples. In fact, the induction of cluster 2 genes was even more robust in WD-fed Setdb2mKO monocytes (Figure 6K-M).

**Figure 6.**
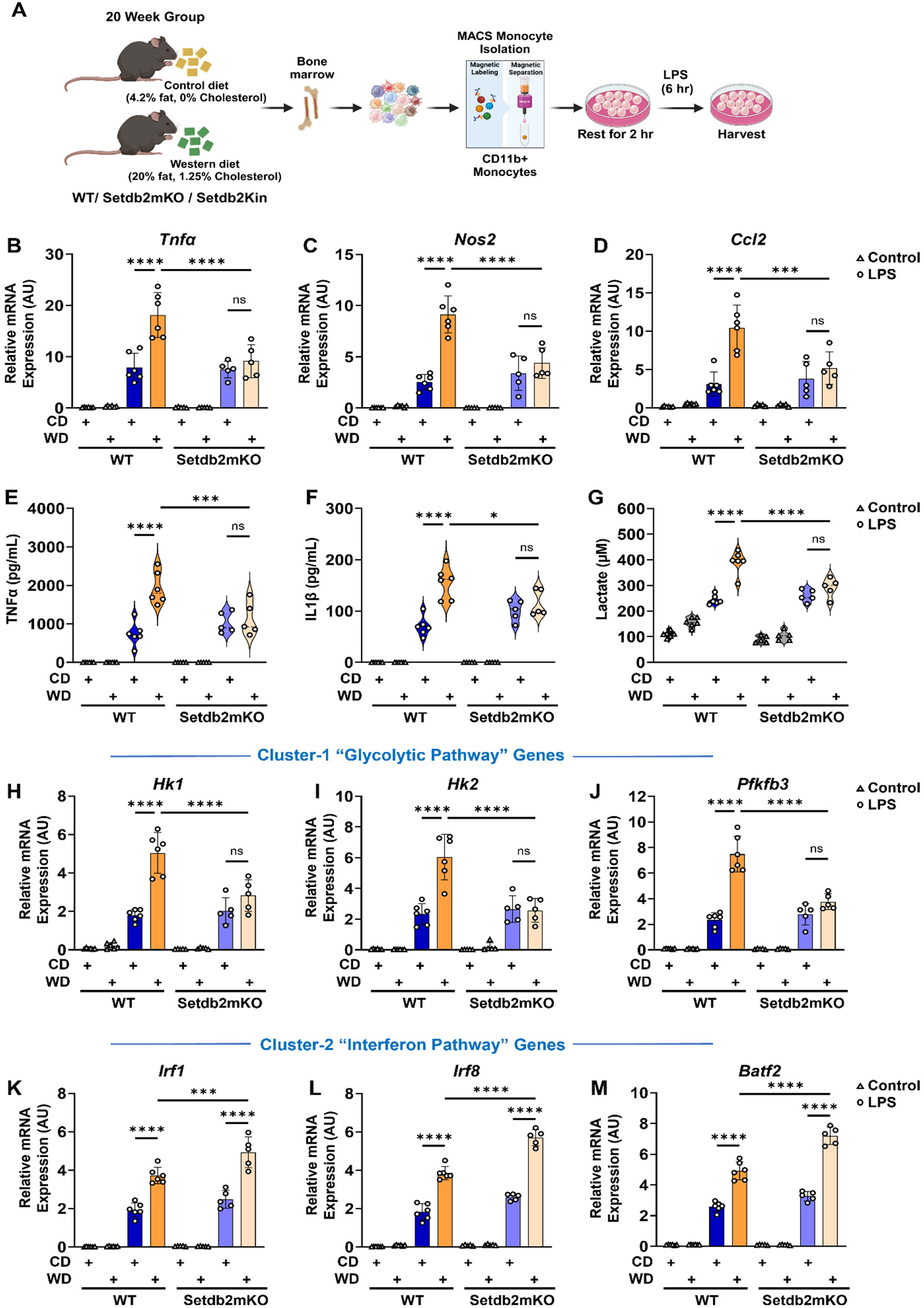
Setdb2 regulates ‘western-diet induced’ innate immune memory of bone-marrow monocytes. (A) Graphical outline showing in vivo western diet-induced training model. CD11b^+^ BM-monocytes were isolated from control-diet or western-diet fed WT or Setdb2mKO mice followed by stimulation with 100ng/mL LPS or culture medium in vitro for 6 h. ("See also Figure S5") (B) mRNA expression of Tnfα (C) Nos2, and (D) Ccl2, (E) Pro-inflammatory cytokine levels of TNFα, and (F) IL-1β, (G) Extracellular lactate levels, (H) mRNA expression of Hk1 (I) Hk2, (J) Pfkfb3 (K) Irf1 (L) Irf8, and (M) Batf2 in control diet, western diet, control diet followed by in vitro LPS restimulation, and western diet followed by in vitro LPS restimulated BM-monocytes isolated from WT and Setdb2mKO mice. Control diet WT (n=6), Control diet Setdb2mKO (n=5), Western diet WT (n=6), and Western diet Setdb2mKO mice (n=5) from one of the three independent experimental group. Data are represented as mean ± SD. ‘n’ represents biological replicates of each strain and treatment type. p-value was calculated using two-way ANOVA with Tukey post-hoc test for multiple comparisons. ∗p < 0.05, ∗∗p < 0.01, and ∗∗∗p < 0.001, ns, not significant change.

The WD feeding response for cluster 1 genes in Setdb2Kin mice was similar to WT (Figure S6A-I) which as the same for β-glucan training in vitro. Additionally, induction of cluster-2 genes by LPS challenge in WD-fed Setdb2Kin monocytes was also similar to WT and Setdb2mKO (Figure S6J-L). These results suggest that regulation of cluster-1 pathway genes by Setdb2 is conserved across pathogenic (β-glucan) and sterile (WD) inflammatory stimuli, whereas cluster-2 pathway gene regulation is differentially affected by these different triggers.

### In vitro training with oxidized-LDL regulates trained immunity responses in bone marrow macrophages similar to ‘Western-diet’ feeding

Western diet feeding is associated with an increased level of oxidized lipids and lipoproteins that contribute to the associated increased chronic inflammation and atherosclerotic burden.^19^ In fact, oxLDL has been shown to substitute for β-glucan as an initial trigger during the two-step in vitro innate immune training challenge.^20^ Thus, we hypothesized that the differential response of cluster-2 genes to WD training above might also be observed for in vitro challenge to oxLDL. Thus, we compared the oxLDL-induced trained immunity in WT and Setdb2mKO mice BM-macrophages. Interestingly, we found that the expression patterns for cluster-1 and cluster-2 associated genes look exactly the same as for the WD feeding results. Here, oxLDL training resulted in super-induction of cluster-1 associated responses in WT, and these effects were blunted in the BM-macrophages isolated from Setdb2mKO mice (Figure 7A-I). However, oxLDL training did not limit the LPS response of cluster-2 pathway gene induction and similar to WD training in vivo cluster 2 genes were also further induced by oxLDL pretraining (Figure 7J-L).

**Figure 7.**
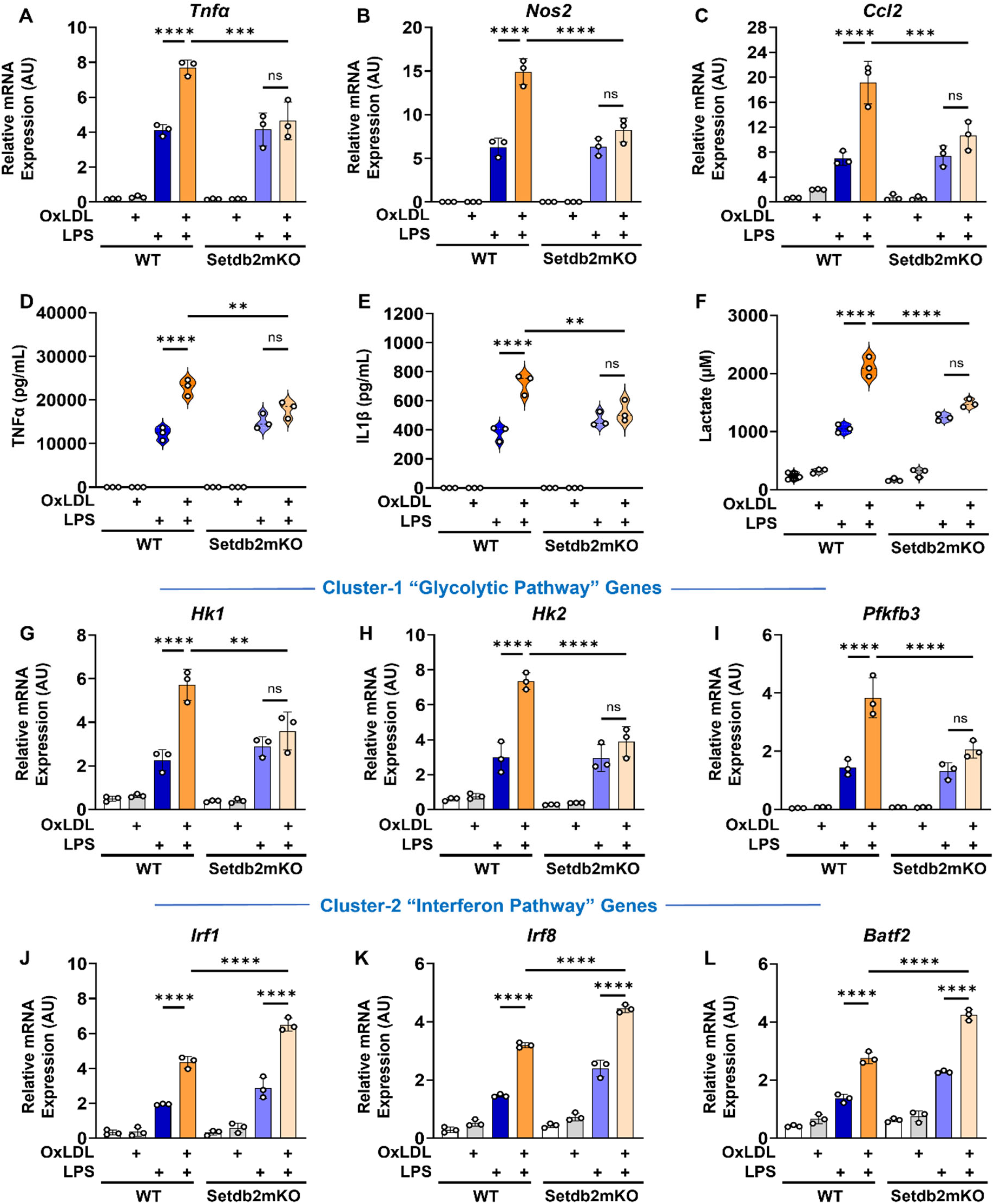
In-vitro training with oxidized-LDL regulates trained immunity responses in bone marrow macrophages similar to ‘Western-diet’ feeding. Bone marrow cells from WT and Setdb2mKO mice were isolated and treated with 10 μg/mL oxLDL (training stimulus) or culture medium for 24 h, cultured for 5 days, and restimulated with 100ng/mL LPS (secondary challenge) or culture medium for 6 h on day 6. Cells were harvested for analysis. (A) mRNA expression of Tnfα, (B) Nos2, and (C) Ccl2, (D) Pro-inflammatory cytokine levels of TNFα, and (E) IL-1β, (F) Extracellular lactate levels, (G) mRNA expression of Hk1 (H) Hk2, (I) Pfkfb3, (J) Irf1 (K) Irf8, and (L) Batf2 in untreated control, oxLDL alone, LPS alone, and oxLDL followed by LPS restimulated BM-macrophages from WT (n = 3) or Setdb2mKO mice (n = 3). Data are represented as mean ± SD. ‘n’ represents biological replicates of each strain and treatment type. p-value was calculated using two-way ANOVA with Tukey post-hoc test for multiple comparisons. ∗p < 0.05, ∗∗p < 0.01, and ∗∗∗p < 0.001, ns, not significant change.

## DISCUSSION

In this study, we identified a dual role for Setdb2 in the innate immune memory response of monocytes/macrophages. When bone marrow-derived cells were treated with either β-glucan or oxidized LDL in vitro as a training agent and then cultured for several days followed by secondary stimulation with LPS on day 6, the cells pre-treated with either training compound displayed a more robust secretion of TNF and IL-1β cytokine in response to the secondary exposure to LPS relative to cells treated only with LPS. There was also enhanced secretion of lactate consistent with the metabolic switch to aerobic glycolysis. Importantly, these responses were accompanied by parallel changes in expression of the corresponding cytokine and glycolytic genes consistent with a transcriptional regulatory mechanism of action. These responses to the two-step training protocol represent the classic measures for the innate immune memory response reported over a decade ago by *Netea and colleagues*.^1,2^ Subsequent studies have demonstrated that the initial stimulus leads in part to chromatin-dependent reprogramming that is established following the training exposure, which can be retained over the course of several days in culture. The reprogramming provides a poised platform to support the more robust response to secondary exposure to LPS. We showed that the training response to LPS was significantly blunted in cells isolated from Setdb2mKO mice. RNA-seq comparisons between WT and Setdb2mKO macrophages revealed 5 unique clusters of gene expression response patterns. Cluster 1 genes were super-induced by LPS in response to β-glucan pre-training, whereas the LPS-mediated induction of cluster 2 genes on day 6 was blunted by β-glucan pre-exposure. Interestingly, both responses were lost in cells from Setdb2mKO mice, consistent with a role for Setdb2 in β-glucan pretraining of genes in both clusters. This differential response uncovered a dual role for Setdb2: stimulating expression of cluster 1 genes while suppressing expression of cluster 2 genes, respectively.

Setdb2 is a member of the KMT1 family of lysine methyltransferases, several of which are known to methylate lysine 9 of histone H3, a modification long known to be associated with inhibition of gene expression. Consistent with this role, others have shown Setdb2 is induced by interferon signaling during the early proinflammatory response in macrophages where it binds to promoters encoding key proinflammatory genes, which are normally silenced later to prevent runaway inflammation. In this context, Setdb2 binding is associated with increased methylation of H3K9, and loss of Setdb2 leads to prolonged stimulation of its target proinflammatory genes coincident with the decrease in promoter-associated lysine 9 methylation. This suggests Setdb2 represses inflammatory gene expression through its lysine methyltransferase function.

In another study, our lab showed that Setdb2 is induced by fasting in the liver, where it interacts with GR to co-activate a subset of GR target genes involved in the hepatic fasting response. In this earlier report, we showed Setdb2 is associated with increased enhancer-promoter looping, gene activation, and reduced H3K9 promoter methylation. Additionally, we showed that a mutant version of Setdb2 that altered key residues involved in binding the methyl-donating SAM cofactor still functioned normally to co-activate GR target genes. These results are consistent with a role for Setdb2 in gene activation that is independent of its potential enzyme activity.

To investigate the catalytic and non-catalytic roles for Setdb2 in the innate training response, we engineered a knock-in mouse model (Setdb2Kin) where two critical amino acids in the SET domain that are critical for SAM binding were changed to alanines. These substitutions were designed to prevent interaction with SAM based on the known structure of the SET domain,^12^ which would eliminate any potential inherent methylation activity. When we repeated the β-glucan training study with Setdb2Kin bone marrow-derived cells, the response to β-glucan training for the super-induction of cluster 1 genes by LPS was maintained, whereas the negative effect on LPS activation of cluster 2 genes was lost. When taken together with the loss of both responses in the Setdb2mKO mouse, these results are consistent with a dual role for Setdb2 in β-glucan-dependent training: a positive role in activation of cluster 1 and a negative role in cluster 2 gene responses.

Based on these two opposite responses and the previous results from our lab and others, we hypothesized that enhancer promoter looping was involved in Setdb2 regulation of cluster 1 gene activation, whereas H3K9 methylation was involved in repression of cluster 2 gene expression. To test these predictions, we utilized 3C to assess enhancer-promoter looping in cluster 1 genes, and we utilized direct ChIP analyses for H3K9 methylation to interrogate cluster 2 gene responses during the two-step β-glucan training experiment. The results showed that cluster 1 and 2 genes were associated with enhancer-promoter looping and H3K9 methylation changes, respectively, consistent with our hypothesis.

This dual role for Setdb2 is similar to the actions of other chromatin modifiers, including G9a and JMJD1A. Similar to Setdb2, G9a is also a member of the KMT1 family of lysine methyltransferases and generates H3K9me2 at target genes to effect gene silencing. However, it also acts as a molecular scaffold factor in the assembly of a larger complex with other coactivators during gene activation where its enzyme activity is not required.^21^ Similarly, JMJD1a is an H3K9 demethylase, and its demethylase activity is involved in the stimulation of white adipose “browning,” but other studies also showed it is involved in enhancer-promoter looping to activate a set of thermogenic target genes in brown adipose tissue, and introducing mutations that eliminate its demethylase activity do not affect its role in chromatin looping.^22^

We also performed ATAC-seq analysis on β-glucan-trained WT vs. Setdb2mKO following the secondary LPS challenge. These studies identified enriched motifs within open chromatin regions associated with the promoters of genes in clusters 1 and 2. As expected, the *NFκB* motif was significantly associated with changing peaks proximal to genes for both clusters. However, the *mafK* motif was only highly associated with peaks close to cluster 2 genes. This is interesting because *mafK* is co-recruited along with the SAM synthesizing *matIIa/b* enzyme complex to chromatin to provide the methyl donor needed for H3K9 methylation in concert with Setdb1.^18^Interestingly, Setdb1 is the only other SET enzyme within the KMT1 family with a bifurcated SET domain.^13^ Further studies are required to determine whether Setdb2 might also be recruited to suppress cluster 2 gene promoters through mafK or a related repressor.

Metabolic disorders, including atherosclerosis, are associated with chronic low-grade inflammation, where the innate immune system contributes to vascular inflammation associated with atherosclerotic plaque formation.^23^ Monocyte infiltration into the arterial wall followed by local responses including hypoxia and accumulation of excess modified lipids results in foam cell formation which accelerates atherosclerosis progression. In fact, the inhibition of monocyte infiltration has protective effects in experimental mouse models of atherosclerosis.^24^ However, it is important to note that the appearance of inflammatory monocytes is not only found within plaques but also in the circulation.^25, 26^ In this regard, a recent study in mice also showed that WD feeding leads to reprogramming of the myeloid cell compartment within the bone marrow, leading to a chronic low-grade state of “sterile inflammation”. It also leads to long-lasting reprogramming of circulating myeloid cells, resulting in a hyper-response to a wide range of different secondary immune stimuli.^4^ These studies suggest that ‘sterile inflammatory’ conditions also induce an innate immune memory response similar to infectious triggers like β-glucan.

In a previous study, we reported that bone marrow transplantation from global Setdb2 deficient mice into WD fed *Ldlr-/-* mice resulted in increased vascular inflammation and aggravated atherosclerosis relative to parallel transplants using bone marrow from control WT mice. In the current study we showed that monocytes isolated from WD-fed WT mice were hyper-responsive to secondary LPS challenge in vitro. We also showed that β-glucan injection into mice also resulted in a similar training response to LPS in macrophages challenged with LPS in vitro. These results are similar to prior reports and are consistent with WD feeding and β-glucan providing an initial signal that effectively marks circulating monocytes similar to the adaptation effect of β-glucan exposure in vitro.

The training response in macrophages isolated from mice after β-glucan injection was similar to the response observed with bone marrow cells treated with β-glucan in vitro, where the induction and repression of cluster 1 and 2 genes was observed, and both responses were lost in the Setdb2mKO mice. However, the results when WD feeding was the source of the training signal were different. The super-induction of cluster 1 genes in response to LPS challenge was abolished in cells from Setdb2mKO, but the inhibition of the LPS response for cluster 2 genes was not observed. In fact, if anything, cluster 2 genes were induced even more robustly by LPS challenge than in cells from WT mice. Oxidized LDL (oxLDL) is generated during WD feeding, and it has been suggested to be a key driver of the chronic inflammatory response associated with atherosclerosis.^19^ In fact, studies showed that in vitro challenge of bone marrow-derived cells with oxLDL elicited a similar training response to β-glucan. When we repeated the in vitro training experiment using oxLDL to challenge WT and Setdb2mKO cells, we observed a very similar response to the in vivo WD training protocol. The super-activation of cluster 1 genes was lost, but the cluster 2 repression response observed in trained WT cells was not observed. In fact, similar to WD training in vivo, cluster 2 genes were stimulated even more robustly by LPS challenge in the oxLDL-pretrained Setdb2mKO cells. Taken together, these results demonstrate that the loss of Setdb2 leads to a bifurcation in the responses elicited by the two different training agents.

Using single-cell RNA-seq on CD45+ cells in the atherosclerosis studies mentioned above, we found that the aggravated atherosclerotic plaques from the mice reconstituted with Setdb2-deficient bone marrow contained significantly fewer T-cells along with significantly enhanced neutrophil infiltration.^17^ A more detailed analysis of our bulk RNAseq data from the β-glucan training experiment reported here showed an enrichment for Ccl chemokine signaling mediators within cluster-1 genes, whereas Cxcl pathway regulators were enriched in cluster 2. Because cluster 1 and 2 genes are uniquely under- and over-represented in the Setdb2mKO, this analysis also reveals a potential underlying mechanism for the differential appearance of T-cell and neutrophil populations in Setdb2-deficient atherosclerotic plaques.

Overall, our evaluation of Setdb2 in the innate immune training response has uncovered an essential role for Setdb2 that involves a previously unknown branching in the downstream pathways from the initial trigger that define the memory response. Our study also reveals that Setdb2 plays a key role in different response patterns to infectious (β-glucan) vs. sterile (WD/oxLDL) inflammatory triggers. Thus, our results not only provide evidence for significant differences in the training response to two different major pro-inflammatory activation pathways, but they also reveal key mechanistic insight into Setdb2’s mode of action for differential transcriptional regulation of the key innate immune response pathways that are affected. This opens up the possibility that targeting Setdb2’s enzymatic vs. scaffold activity may result in context-dependent potential therapeutic targeting of the divergent pathways in inflammatory or metabolic disorders.

## RESOURCE AVAILABILITY

### Lead contact

Further information and requests for resources and reagents should be directed to and will be fulfilled by the lead contact, Timothy Osborne.

### Materials availability

All unique reagents generated in this study are available from the lead contact with a completed material transfer agreement.

### Data and code availability

Sequencing data sets performed in this study are available at the NCBI GEO under accession numbers: GSE290872

## ACKNOWLEDGMENTS

The authors express their appreciation to Professor Andreas Bergthaler for supplying the Setdb2 antibody. Authors recognize Anne Lynch for her critical role in maintaining the mouse colony and performing genotyping. Gratitude is extended to Crystal Young-Erdos for her help with ChIP analysis, as well as to Bence Daniel and Kandarp Joshi for their helps to 3C-qPCR analysis. Yang Yang and members of the Osborne lab are thanked for their valuable scientific discussions and feedback. The institute’s vivarium and Shared Resources staff are acknowledged for ensuring a healthy environment for the mouse colony and maintaining key equipment. Sequencing was performed at the Genetic Resources Core Facility, RRID* SCR_018669, within the Johns Hopkins Department of Genetic Medicine, Baltimore, MD. Schematic images were created using BioRender software. This work was supported by National Institutes of Health grant R01 DK124343 to T.O.

## AUTHOR CONTRIBUTIONS

Conceptualization, T.O. and A.J.; A.J. performed the experiments. L.H. designed the bioinformatic approaches and carried out the analyses. D.L.W., provided β-glucan and related scientific input. A.J. writing—original draft, A.J., L.H. and T.O.; writing—review & editing. T.O. obtained funding and provided supervision. All authors provided input and approved the final version of the manuscript.

## DECLARATION OF GENERATIVE AI AND AI-ASSISTED TECHNOLOGIES

During the preparation of this work, the authors have not used any AI-assisted technology or generative images, and takes full responsibility for the content of the publication.

## SUPPLEMENTAL INFORMATION

**Document S1. Figures S1–S6 and Table S1**

## KEY RESOURCES TABLE

| REAGENT OR RESOURCE | SOURCE | IDENTIFIER |
| --- | --- | --- |
| **Antibodies** |  |  |
| Anti-Setdb2 | Professor Andreas Bergthaler, Molecular Immunology, MedUni Vienna, Austria | N/A |
| Anti-YY1 | Cell Signaling Technology | Cat# 46395, RRID:AB_2799302 |
| Anti-H3K9me3 | Abcam | Cat# ab8898, RRID:AB_306848 |
| Anti-H3K9me2 | Abcam | Cat# ab176882, RRID:AB_2895140 |
| Anti-H3K9me1 | Abcam | Cat# ab176880, RRID:AB_2751009 |
| Anti-H3 | Cell Signaling Technology | Cat# 14269, RRID:AB_2756816 |
| Anti-β-Actin | Cell Signaling Technology | Cat# 3700, RRID:AB_2242334 |
| Rabbit IgG, polyclonal-Isotype Control | Abcam | Cat# ab171870, RRID:AB_2687657 |
| Goat Anti-Mouse IgG H&L (HRP) | Abcam | Cat# ab6789, RRID:AB_955439 |
| **Bacterial and virus strains** |  |  |
| **Biological samples** |  |  |
| **Chemicals, peptides, and recombinant proteins** |  |  |
| β-glucan (b1,3-(D)-glucan) | Professor David Williams, College of Medicine, Johnson City, USA | N/A |
| Murine GM-CSF | PeproTech | Cat# 315-03 |
| Lipopolysaccharides (LPS) | Sigma-Aldrich | Cat# L7770 |
| Oxidized Low Density Lipoprotein from Human Plasma (OxLDL) | Thermo Fisher Scientific | Cat# L34357 |
| Dulbecco's Modification of Eagle's Medium (DMEM) | Corning | Cat# 10-013-CV |
| RPMI 1640 Medium (Dutch modification) | Thermo Fisher Scientific | Cat# 22409015 |
| Gentamicin | Thermo Fisher Scientific | Cat# 15750060 |
| Glutamax | Thermo Fisher Scientific | Cat# 35050061 |
| Pyruvate | Thermo Fisher Scientific | Cat# 11360070 |
| Fetal Bovine Serum (FBS) | Corning | Cat# 35-011-CV |
| Penicillin/Streptomycin | Thermo Fisher Scientific | Cat# 15-140-122 |
| ACK-RBC lysis buffer | Neta Scientific | Cat# 118-156-101 |
| Protease inhibitor cocktail | Milipore Sigma | Cat# 11836153001 |
| PVDF membranes | Thermo Fisher Scientific | Cat# IPVH00005 |
| Any kD™ Mini-PROTEAN | Bio-Rad | Cat# 4569034 |
| SYBR green master mix | Quantabio | Cat# 101414-152 |
| TaqMan Fast Advanced PCR Master Mix | Applied Biosystems | Cat# 44-445-57 |
| NotI-restriction endonuclease | New England BioLabs (NEB) | Cat# R0189L |
| MboI-restriction endonuclease | New England BioLabs (NEB) | Cat# R0147L |
| Disuccinimidyl glutarate (DSG) | ProteoChem | Cat# C1104 |
| Formaldehyde | Thermo Fisher Scientific | Cat# PI28908 |
| Protein G Dynabeads | Invitrogen | Cat# 10004D |
| **Critical commercial assays** |  |  |
| Cholesterol assay kit | Cell Biolabs, Inc | Cat# STA-384 |
| TNF-alpha Quantikine ELISA Kit | R&D Systems | Cat# MTA00B |
| IL-1β Quantikine ELISA Kit | R&D Systems | Cat# MLB00C |
| L-lactate assay kit | Sigma-Aldrich | Cat# MAK329 |
| Monocyte isolation kit | Miltenyi | Cat# 130-100-629 |
| BCA Protein Assay Kit | Thermo Fisher Scientific | Cat# A55864 |
| Super Signal™ West Femto Maximum Sensitivity Substrate ECL-Kit | Thermo Fisher Scientific | Cat# 34095 |
| RNeasy kit for RNA purification | Qiagen | Cat# 74106 |
| High-Capacity cDNA Reverse Transcription Kit | Applied Biosystems | Cat# 4368814 |
| NEBNext® Poly(A) mRNA Magnetic Isolation Module | New England BioLabs (NEB) | Cat# E7490S |
| NEBNext Ultra II Directional RNA Library Prep Kit for Illumina | New England BioLabs (NEB) | Cat# E7760S |
| MinElute PCR purification kit | Qiagen | Cat# 28004 |
| Tn5 transposase | Illumina | Cat# 20034197 |
| CutSmart Digestion buffer | New England BioLabs (NEB) | Cat# B6004S |
| Serum-separator tube | Thermo Fisher Scientific | Cat# BD 365967 |
| **Deposited data** |  |  |
| Bulk RNA-sequencing | This paper | GSE290872 |
| ATAC- sequencing | This paper | GSE290872 |
| **Experimental models: Cell lines** |  |  |
| Mouse bone-marrow macrophages | C57BL/6J, LysM<sup>cre+</sup>, Setdb2mKO, Setdb2Kin | N/A |
| Mouse peritoneal macrophage | LysM<sup>cre+</sup>, Setdb2mKO | N/A |
| Mouse bone marrow CD11b<sup>+</sup> monocyte | C57BL/6J, LysM<sup>cre+</sup>, Setdb2mKO, Setdb2Kin | N/A |
| **Experimental models: Organisms/strains** |  |  |
| C57BL/6J mice | The Jackson Laboratory | N/A |
| LysM<sup>cre+</sup> mice | The Jackson Laboratory | N/A |
| Setdb2<sup>flox/flox</sup> LysM<sup>cre+</sup> mice (Setdb2mKO) | This paper | N/A |
| Setdb2 Knock-in mice (Setdb2Kin) | This paper | N/A |
| **Oligonucleotides** |  |  |
| Primer for mRNA expression-qPCR | This paper | see Table S1 |
| TaqMan probe and primer for 3C-qPCR | This paper | see Table S1 |
| Primer for ChIP-qPCR | This paper | see Table S1 |
| Primer for sanger-seq | This paper | see Table S1 |
| **Recombinant DNA** |  |  |
| **Software and algorithms** |  |  |
| nf-core/rnaseq v3.0 | (Ewels et al., 2020)<sup>27</sup> | https://github.com/nf-core/rnaseq |
| nf-core/atacseq v2.0 | (Ewels et al., 2020)<sup>27</sup> | https://github.com/nf-core/atacseq |
| DeepTools v3.5.1 | (Ramirez et al., 2016)<sup>28</sup> | https://github.com/deeptools/deepTools/tree/develop |
| FastQC v0.11.9 | – | https://www.bioinformatics.babraham.ac.uk/projects/fastqc/ |
| bamCoverage | (Ramirez et al., 2016)<sup>28</sup> |  |
| STAR | (Dobin et al., 2013)<sup>29</sup> | https://github.com/alexdobin/STAR |
| Salmon | (Patro et al., 2017)<sup>30</sup> | https://github.com/COMBINE-lab/salmon |
| DiffBind v3.6.3 | (Ross-Innes et al., 2012)<sup>31</sup> | https://bioconductor.org/packages/release/bioc/html/DiffBind.html |
| EdgeR v3.38.4 | (Robinson et al., 2010)<sup>32</sup> | https://bioconductor.org/packages/release/bioc/html/edgeR.html |
| IGV v2.14.0 | (Robinson et al., 2011)<sup>33</sup> | https://software.broadinstitute.org/software/igv/ |
| ComplexHeatmap | (Gu et al., 2016)<sup>34</sup> |  |
| EnrichR | (Kuleshov et al., 2016)<sup>35</sup> |  |
| HOMER v4.11 | (Heinz et al., 2010)<sup>36</sup> | http://homer.ucsd.edu/homer/index.html |
| chromVAR | (Schep et al., 2017)<sup>37</sup> |  |
| Picard | – | http://broadinstitute.github.io/picard/ |
| Bowtie2 | (Langmead and Salzberg, 2012)<sup>38</sup> |  |
| GraphPad Prism 10 |  |  |
| BWA v0.7.17 | (Li et al., 2009)<sup>39</sup> | https://github.com/lh3/bwa |
| BEDTools v2.30.0 | (Quinlan and Hall, 2010)<sup>40</sup> | https://bedtools.readthedocs.io/en/latest/ |
| Samtools v1.16 | (Li et al., 2009)<sup>39</sup> | http://www.htslib.org/ |
| MACS2 v2.2.7.1 | (Zhang et al., 2008)<sup>41</sup> | https://pypi.org/project/MACS2/ |
| DESeq2 v1.36.0 | (Love et al., 2014)<sup>42</sup> | https://bioconductor.org/packages/release/bioc/html/DESeq2.html |
| ENCODE mm10 blacklist | (Amemiya et al., 2019)<sup>43</sup> | https://github.com/Boyle-Lab/Blacklist |
| **Other** |  |  |
| Western diet | Research Diets, Inc | Cat# D12108Ci |
| Control diet | Research Diets, Inc | Cat# D12102Ci |
| Bacterial artificial chromosome (BAC) clone spanning *Tnf* locus | CHORI Bac Pac Resource Center | Clone RP23-446C22 |
| Bacterial artificial chromosome (BAC) clone spanning *Ccl2* locus | CHORI Bac Pac Resource Center | Clone RP23-230M24 |

## EXPERIMENTAL MODEL AND STUDY PARTICIPANT DETAILS

### Mice

All animal experiments were carried out in compliance with ethical guidelines approved by the Institutional Animal Care and Use Committee (IACUC) of Johns Hopkins University (license number MO24C317). Mouse colonies were handled by ’Charles River Laboratories’ according to the institute’s animal facility’s regulatory standards at Johns Hopkins All Children’s Hospital.

*C57BL/6J* and *lysozyme-Cre (Lyz2-cre)+* mice were purchased from The Jackson Laboratory and maintained in-house. *Setdb2flox* conditional mice with *a C57BL/6J* background were generated using embryonic stem (ES) cells derived from gene-trap mice as described previously.^16^ Myeloid-specific Setdb2-deficient strain (Setdb2mKO) was generated by crossing *Setdb2^flox/flox^* mice with the *lysozyme-Cre (Lyz2-cre)+* animals and confirmed through genotyping and qPCR. The Global Setdb2 knock-in (Setdb2Kin) mouse line was generated by introducing two germline mutations in the SET domain of the Setdb2 gene using gene targeting technology by the Ingenious Targeting Laboratory. The specificity of the Setdb2 knock-in mutations was further confirmed by Sanger sequencing (Supplemental Figure S2). Eight-to twelve-week-old female mice were used for in vivo and in vitro experiments with at least 3 mice per genotype. Age-matched *lysozyme-Cre (Lyz2-cre)+* and wild-type *C57BL/6J* mice were used as a control for Setdb2mKO and Setdb2Kin mice, respectively. All mice had an *ad libitum* supply of food and water throughout housing under a 12-hour light-dark cycle in a helicobacter/norovirus-free room. Unless otherwise stated, mice were sacrificed within one hour of the dark/light transition.

### Bone-marrow macrophage isolation and in vitro trained immunity model

Bone marrow (BM) cells were isolated as previously described.^44^ Isolated bone marrow cells were washed and resuspended in DMEM culture medium supplemented with 10% FBS (35-011-CV, Corning) and 1% penicillin-streptomycin (15-140-122, Thermo Fisher Scientific). Cells were counted on a Countess cell counter instrument (Invitrogen), and 1×10^6^ cells were plated into each well of a flat-bottom 6-well plate with 20 ng/ml GM-CSF (315-03, PeproTech). Cells were kept in a 5% CO₂ incubator at 37°C for 1 hour. After that, cells were treated with 5 µg/ml β-glucan (provided by Professor David Williams) or 10 µg/mL oxLDL (L34357, Thermo Scientific) for 24 hours as a training component. Cells were washed once on the following day and then maintained in complete DMEM media supplemented with 20 ng/ml GM-CSF for 5 days. The medium was changed on the 3rd day of the incubation. On day 6, cells were washed and re-stimulated with culture medium alone or medium supplemented with 100 ng/mL LPS (L7770, Sigma-Aldrich) and cultured for 6 hours. After that, cell supernatants were collected and stored at −20°C for later measurement of lactate and cytokine levels, and the cells were washed and collected for further analysis.

### In vivo β-glucan training and peritoneal macrophage isolation

In vivo, β-glucan training was assessed by intraperitoneal administration of 1 mg β-glucan in 150 μL of endotoxin-free phosphate-buffered saline (PBS) into 8-week-old female Setdb2mKO or wild-type (*Lyz2-cre)+* mice. Control mice were injected with 150 ul PBS alone. On day 6, mice were euthanized, and peritoneal macrophages were isolated as described.^45^ Briefly, the peritoneum was flushed multiple times with peritoneal macrophage isolation media (3% FBS in 1x PBS). The peritoneal fluid was centrifuged at 500 g for 10 min, and the RBCs in the pelleted cells were lysed by resuspending the cell pellet into RBC lysis buffer (118-156-101, Neta Scientific). Cells were again collected by centrifugation and resuspended in complete RPMI medium (22409015, Thermo Scientific) supplemented with 10 μg/mL gentamicin (15750060, Thermo Scientific), 2 mM Glutamax (35050061, Thermo Scientific), and 1 mM pyruvate (11360070, Thermo Scientific). 5 x 10^5^ cells were plated per well in a 24-well flat-bottom plate. After 2 hours, suspension cells were removed by washing with a warm complete RPMI medium, and adherent macrophages were treated with 100 ng/mL LPS (L7770, Sigma-Aldrich) as a secondary stimulus. After 6 hours of treatment, cell supernatants were collected and stored at −20°C for cytokine and lactate measurement, and cells were washed and processed for RNA extraction and qPCR analysis.

### In vivo western diet feeding-induced hyperlipidemia mice model

To develop western diet-induced hyperlipidemia/hypercholesterolemia in mice, Setdb2mKO, Setdb2Kin, and wild-type mice were fed with a Western diet (D12108Ci, Research Diets, Inc.) containing 1.25% cholesterol, 20% fat, 22.6% protein, and 45.3% carbohydrate or a control diet (D12102Ci, Research Diets, Inc.) consisting of 4.2% fat, 22.6% protein, and 66.3% carbohydrates with no cholesterol for 20 weeks. Mice were constantly monitored for their health. Body weights were measured every two weeks throughout the experiment. Diets were frequently replenished into the cage to avoid excessive oxidation of the fat-rich diet. After 20 weeks, mice were euthanized, and blood, tissue, and bone marrow were collected for further analyses.

## METHOD DETAILS

### Cholesterol measurement

After 20 weeks of western diet feeding, mice were euthanized, and blood was collected from the left ventricle via cardiac puncture. Blood samples were collected in a serum-separating tube (BD 365967, Fisher Scientific) containing clot activator gel. The serum was separated by centrifugation at 2000 g for 15 minutes at 4°C. Total cholesterol was measured in serum using a colorimetric cholesterol assay kit (STA-384, Cell Biolabs, Inc.), as per the manufacturer’s protocol. Plates were analyzed on the BioTek Cytation Imaging and Microplate Reader (Agilent).

### Bone marrow (BM)-monocyte isolation from Western diet-induced hyperlipidemia mice

To study western diet-induced innate immune training, CD11b+ BM-monocytes were isolated from bone marrow via magnetic (MACS) sorting using a monocyte isolation kit (Miltenyi, 130-100-629). Non-target cells were depleted using a cocktail of biotin-conjugated monoclonal antibodies, and CD11b+ monocytes were separated with 95% purity. Cells were resuspended in complete RPMI medium supplemented with 10 μg/mL gentamicin, 2 mM Glutamax, and 1 mM pyruvate. 3 x 10^5^ cells were plated per well of a 48-well plate. After 1-hour incubation in a 5% CO₂ incubator at 37°C, cells were stimulated with LPS as a secondary challenge. After 6 hours, cell supernatants were collected and stored at −20°C for cytokine and lactate measurement, and cells were washed and processed for RNA extraction and qPCR analysis.

### Cytokine production and lactate measurement

Cell culture supernatants were collected and centrifuged at 2000 g for 10 min at 4°C to remove any cell debris. Cell supernatants were separated and stored at -20°C until further analysis. Cytokine levels were measured using commercially available enzyme-linked immunosorbent assay (ELISA) kits for TNFα (MTA00B, R&D Systems) and IL-1β (MLB00C, R&D Systems), as per the manufacturer’s protocol. Extracellular lactate production was measured in cell supernatant using the L-lactate assay kit (MAK329, Sigma-Aldrich) according to the manufacturer’s instructions. Plates were analyzed using BioTek Cytation Imaging and Microplate Reader (Agilent).

### Immunoblot

BM-macrophages from Setdb2mKO and WT mice were collected by centrifugation at 500 g at 4°C for 10 minutes. BM-macrophages were then lysed by resuspending the cell pellet in lysis buffer (50 mM Tris, 0.1% MEA, 1 mM EDTA, 1 mM PMSF, 0.5% Triton X-100) containing 1x protease inhibitor cocktail (11836153001, Milipore Sigma). The cell-lysis buffer mixture was incubated on ice for 20 min with intermixing at 5-minute intervals to ensure complete lysis. The lysed cells were centrifuged at 13,000 g at 4°C for 15 min. The protein concentration in the supernatant was measured with the BCA Protein Assay Kit (A55864, Thermo Scientific) at a wavelength of 562 nm using a BioTek Cytation Imaging and Microplate Reader (Agilent). The protein samples were normalized with a BSA standard (stock: 2 mg/mL, Thermo Scientific). The samples were then diluted up to 1 mg/mL concentration with lysis buffer, and 80 µg of protein for each sample was mixed with 4x SDS denaturation buffer (0.125 M Tris-HCl, 20% glycerol, 4% SDS, 0.02% bromophenol blue dye, 10% MEA, pH 6.8). Samples were heat denatured by incubating at 95°C for 10 min and loaded onto Any kD™ Mini-PROTEAN (4569034, Bio-Rad) SDS-polyacrylamide gels, and after electrophoresis, the gels were transferred onto PVDF membranes (IPVH00005, Thermo Fisher) using the wet-transfer blotting method at a constant 30 volt current overnight in a cold room. Membranes were then blocked with 5% non-fat dry milk in Tris-buffered saline with Tween 20 (TBST) for 2 hours at room temperature. After blocking, membranes were washed once and incubated with primary antibodies for Setdb2 (monoclonal antibody, a gift from Professor Andreas Bergthaler, Austria) or YY1 (46395, Cell Signaling) diluted in 2.5% BSA in 1xTBST with 1:500 or 1:3000 dilution, respectively, at 4°C overnight. Membranes were washed three times with 1xTBST for 15 min at RT and incubated with horseradish peroxidase-labeled anti-mouse secondary antibodies (ab6789, Abcam) at a 1:10000 dilution with 5% non-fat dry milk in TBST for 2 hours at RT. The targeted protein bands were visualized using Super Signal™ West Femto Maximum Sensitivity Substrate ECL-Kit (34095, Thermo Scientific).

### Real-time quantitative PCR

Total RNA was isolated from cells using the RNeasy kit (74106, Qiagen) according to the manufacturer’s instructions. The quality of the RNA was assessed on a Nanodrop 8000 Spectrophotometer (Thermo Scientific). Total RNA was reverse transcribed using a High-Capacity cDNA Reverse Transcription Kit (4368814, Applied Biosystems) as per the manufacturer’s protocol. Quantitative real-time PCR (qPCR) reactions were performed on the CFX Opus 384 Real-Time PCR system (Bio-Rad) using SYBR green master mix (101414-152, Quantabio). Relative mRNA quantities were calculated using the comparative threshold cycle (CT) values. mRNA transcripts were normalized to the L32 housekeeping gene. The primer sequences used for qPCR are available in Table S1.

### RNA-sequencing

Total RNA was isolated from WT and Setdb2mKO BM-macrophages using the RNeasy kit (Qiagen) according to the manufacturer’s instructions. The quality of the RNA was measured on an Agilent BioAnalyzer. High-quality RNA samples were submitted to the Genetic Resource Core Facility (GRCF) at Johns Hopkins University for cDNA library preparation and sequencing. Briefly, mRNAs were enriched by removing ribosomal RNA using NEBNext® Poly(A) mRNA Magnetic Isolation Module (E7490S, New England BioLabs) as per the manufacturer’s instructions. RNA-Seq libraries were prepared using a NEBNext Ultra II Directional RNA Library Prep Kit for Illumina (E7760S, New England BioLabs). Sequencing was performed on a Novaseq-6000 instrument (Illumina). 40-50 million paired-end reads were obtained for each sample, and at least three biological replicates were used from all treatment groups for sequencing.

### Chromatin accessibility profiling by ATAC-sequencing

BM-macrophages from WT and Setdb2mKO mice were scraped from the plates and counted. 60,000 cells per treatment condition were used for each sample. Cells were washed with ice-cold PBS, and nuclei were isolated using lysis buffer (1% Triton-X-100, 150 mM NaCl, 0.1% SDS, 1 mM EDTA, and 20 mM Tris-HCl pH 8.0). Tagmentation was performed in nuclei, using Tn5 transposase in a reaction mixture containing 10.5 μl H₂O, 12.5 μl 2x tagmentation buffer, and 2 μl Tn5 transposase (20034197, Illumina) at 37°C for 30 minutes. The tagmentation reaction was stopped by adding 75 μl TE buffer. DNA was further isolated using Qiagen Minelute columns and amplified by PCR using Nextera primers. PCR reaction clean-up was performed by adding 90 μl of AMPure XP beads to a 50 μl PCR reaction. Samples were submitted to the Genetic Resource Core Facility (GRCF) at Johns Hopkins University for the following fragment analysis by bioanalyzer and sequencing. Libraries were sequenced to a depth of 40-50 million paired-end reads on a Novaseq-6000 instrument (Illumina).

### Chromosome Conformation Capture (3C) qPCR

3C-qPCR analysis was performed in WT and Setdb2mKO BM-macrophages following a previously published protocol ^16,46^ with some modifications. Briefly, 10^6^ BM-macrophages were crosslinked with 1% formaldehyde for 10 min, followed by quenching with 125 mM glycine for 5 min. Nuclei were isolated with a buffer containing 0.1% SDS, 1 mM EDTA, 1% Triton x-100, 150 mM NaCl, and 20 mM Tris, pH 8.0, on rotation at 4°C for 1 hour. Nuclei were then lysed through CutSmart Digestion buffer (New England Biolabs, NEB) with 0.5% SDS at 62°C for 10 minutes. Reactions were terminated by adding 10% Triton-X followed by incubation at 37°C for 15 minutes. The chromatin was then digested with 8 U of MboI (R0147L, New England Biolabs) restriction endonuclease overnight at 37°C. After overnight digestion, chromatin was subjected to proximity ligation using NEB ligation buffer containing 1% Triton-X100 with a total volume of 1.2 mL per reaction for 6-hour incubation at RT with constant rotation. Ligated nuclei were then subjected to Proteinase K digestion, followed by decrosslinking with 5M NaCl at 68°C overnight. DNA was purified using Minelute columns (Qiagen). The resulting 3C template DNA was adjusted to a concentration of 50 ng/uL. GAPDH promoter sequences were used as a reference, and TaqMan real-time quantitative PCR was used to assess target promoter-enhancer interactions. Potential enhancer sequences were selected from the ATACseq profiling data and the epilogos database. Bacterial artificial chromosome (BAC) clones from the CHORI BacPac Resource Center spanning the *Tnf* (clone RP23-446C22), *Ccl2* (clone RP23-230M24), and *Hk2* (clone RP23-346H3) loci were used to make control libraries for normalizing relative PCR amplification efficiencies. BAC clones were processed similarly as 3C template DNA with one additional digestion step with NotI (R0189L, New England Biolabs) endonuclease to linearize the DNA followed by purification. TaqMan probe and primer sequences are listed in the supplemental Table S1.

### Chromatin Immunoprecipitation (ChIP-qPCR)

ChIP-qPCR was performed as previously described.^16,47^ 5×10^6^ BM-macrophages were crosslinked with 50 mM disuccinimidyl glutarate (DSG, C1104 - ProteoChem) for 30 minutes, subsequently with 10 minutes of incubation with 1% formaldehyde. Glycine was added to quench formaldehyde, and nuclei were isolated using ChIP lysis buffer containing 1% Triton-X-100, 150 mM NaCl, 0.1% SDS, 1 mM EDTA, and 20 mM Tris-HCl pH 8.0. Nuclei were then sheared using a Bioruptor 300 instrument (Diagenode) with the following specification: output 4, duty cycle 60%, 15 sec X 6 times, and DNA fragments of ∼2 kb were achieved. Excess shearing was avoided as it decreases H3K9me3 ChIP efficiency.^48^ 50 µg of sheared chromatin was used for overnight immunoprecipitation with the following antibodies: H3K9me3 (ab8898, Abcam), H3K9me2 (ab176882, Abcam), H3K9me1 (ab176880, Abcam), total H3 (14269S, Cell Signaling), and IgG (ab171870, Abcam). Antibody chromatin complexes were pulled down with Protein G Dynabeads (10004D, Invitrogen), followed by washing once with IP wash buffer-1 (0.1% NaDOC, 1% Triton-X-100, 500 mM NaCl, 0.1% SDS, 1 mM EDTA, and 20 mM Tris pH 8.0), once with IP wash buffer-3 (0.5% NaDOC, 0.5% NP-40, 0.25 M LiCl, 1 mM EDTA, and 20 mM Tris pH 8.0), and lastly, once with TE buffer (200 mM Tris, pH 8.0 and 10 mM EDTA). Chromatin was then separated from the beads with elution buffer containing 1% SDS and 100 mM NaHCO₃ through vigorous shaking for 20 minutes, followed by decrosslinking by overnight incubation with 5 M NaCl at 65°C. Immunoprecipitated DNA was further processed for qPCR analysis using a MinElute PCR purification kit (Qiagen) and quantified by Qubit. Primer sequences used for the ChIP-qPCR are listed in Table S1. qPCR reactions were performed using SYBR green master mix (101414-152, Quantabio) on the CFX Opus 384 Real-Time PCR system (Bio-Rad). Samples were normalized with their input control, and H3K9me3, H3K9me2, and H3K9me1 were further normalized with total H3-enriched signals.

## QUANTIFICATION AND STATISTICAL ANALYSIS

### RNA-seq analysis and gene expression data quantification

The paired-ended mRNA sequence reads were analyzed using the nf-core/RNAseq v3.0 pipeline.^27^ Briefly, FastQC (https://github.com/s-andrews/FastQC) was used to quality-check the raw paired-end reads and align them with the *mm10* (GRCm38) genome assembly using default parameters of STAR.^29^ Gene quantification was performed using Salmon.^30^ *DeepTools* and *bamCoverage* ^28^ were used to generate normalized coverage density tracks (bigwig files). Genes containing <10 CPM were separated, and only protein-coding genes were used for further analysis. False discovery rate (FDR) < 0.05 was considered a statistically significant difference calculated by the GLM test using R package *edgeR.*^32^ Multidimensional scaling on the normalized values and hierarchical clustering were performed and visualized in R. *k-means* and hierarchical clustering on the scaled CPM values (k=5) were performed using the Complex Heatmap package.^34^

### GO and GSEA

We performed GSEA using Enrichr.^35^ Hallmark gene sets of MSigDB were utilized and kept significantly enriched terms with P < 0.05.^49, 50, 51^

### ATAC-seq analysis

ATAC-seq paired-end sequencing reads were processed using the nf-core/atacseq pipeline (version 2.0) with default settings. Quality control (QC) was performed using FastQC and Trim Galore for adapter trimming and read filtering. Reads were aligned to the mm10 reference genome using Bowtie2, and duplicate reads were marked or removed using Picard. The peaks representing open chromatin regions were identified with MACS2 and annotated to genomic features using HOMER. Coverage tracks (bigwigs) were created with DeepTools. Differential chromatin accessibility was performed using DiffBind (filter = 3, min overlap = 2, summits = 75).^31^

### Transcription factor activity analysis

The analysis of transcription factor (TF) activity was performed using chromVAR.^37^ Count matrices of accessible regions were generated from ATAC-seq data and used as input. chromVAR was applied to compute deviations in accessibility across samples, using a set of mouse transcription factor motif annotations from CisBP.^52^ Data were first normalized to account for differences in sequencing depth and GC content. Motif accessibility scores (deviation z-scores) were calculated to identify variability in TF binding across conditions. The results were used to infer changes in regulatory activity and highlight condition-specific transcription factor dynamics.

### Statistical analysis

Statistical significance was determined using two-way ANOVA with a Tukey post-hoc test for multiple comparisons. Statistical parameters such as the exact value of n (number of biological replicates), the center, dispersion, and precision measures (mean ± SD) and statistical significance P-value < 0.05 represented with asterisks are described in the figure legends. In bar graphs, error bars represent SD (standard deviation), and bars denote the mean of the indicated number of biological replicates. Quantification and statistical analysis for RNA-seq and ATAC-seq data are explained in detail in the above methods section. Two-way ANOVA analysis was performed using GraphPad Prism 10 (GraphPad Software) with 95% confidence intervals. P-value < 0.05 was considered statistically significant (*, P < 0.05; **P < 0.01; ***, P < 0.001; ****, P < 0.0001).

