## Supplemental material for "Setdb2 Regulates Inflammatory Trigger-Induced Trained Immunity of Macrophages Through Two Different Epigenetic Mechanisms"

**This file includes Supplemental figures S1-S6, figure legends and Table S1**

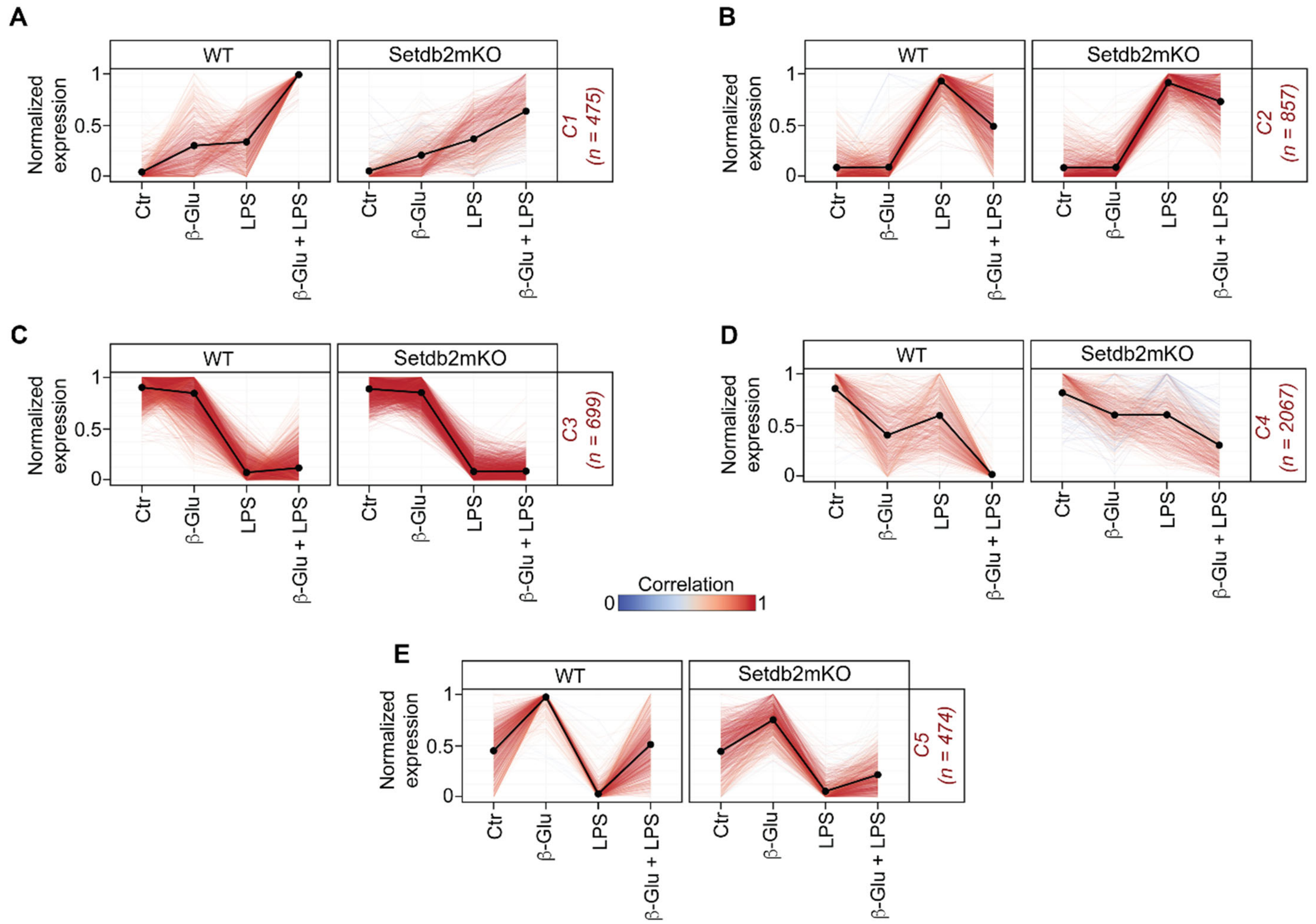

**Figure S1. Hierarchical clustering analysis in β-glucan-induced trained immunity of BM-macrophages (Related to Figure 2)**
(A-E) Line plots showing k-means (k = 5) and hierarchical clustering (n = 3) for genes expressed in untreated control, β-glucan alone, LPS alone, and β-glucan followed by LPS restimulated BM-macrophages from WT (n = 3) and Setdb2mKO mice (n = 3).

Statistical analysis for RNA-seq data was performed using DESeq2 with its default settings. Data are represented as mean ± SD. 'n' represents biological replicates of each strain and treatment type. p-value was calculated using two-way ANOVA with Tukey post-hoc test for multiple comparisons. $*p < 0.05$, $**p < 0.01$, and $***p < 0.001$, ns, not significant change.

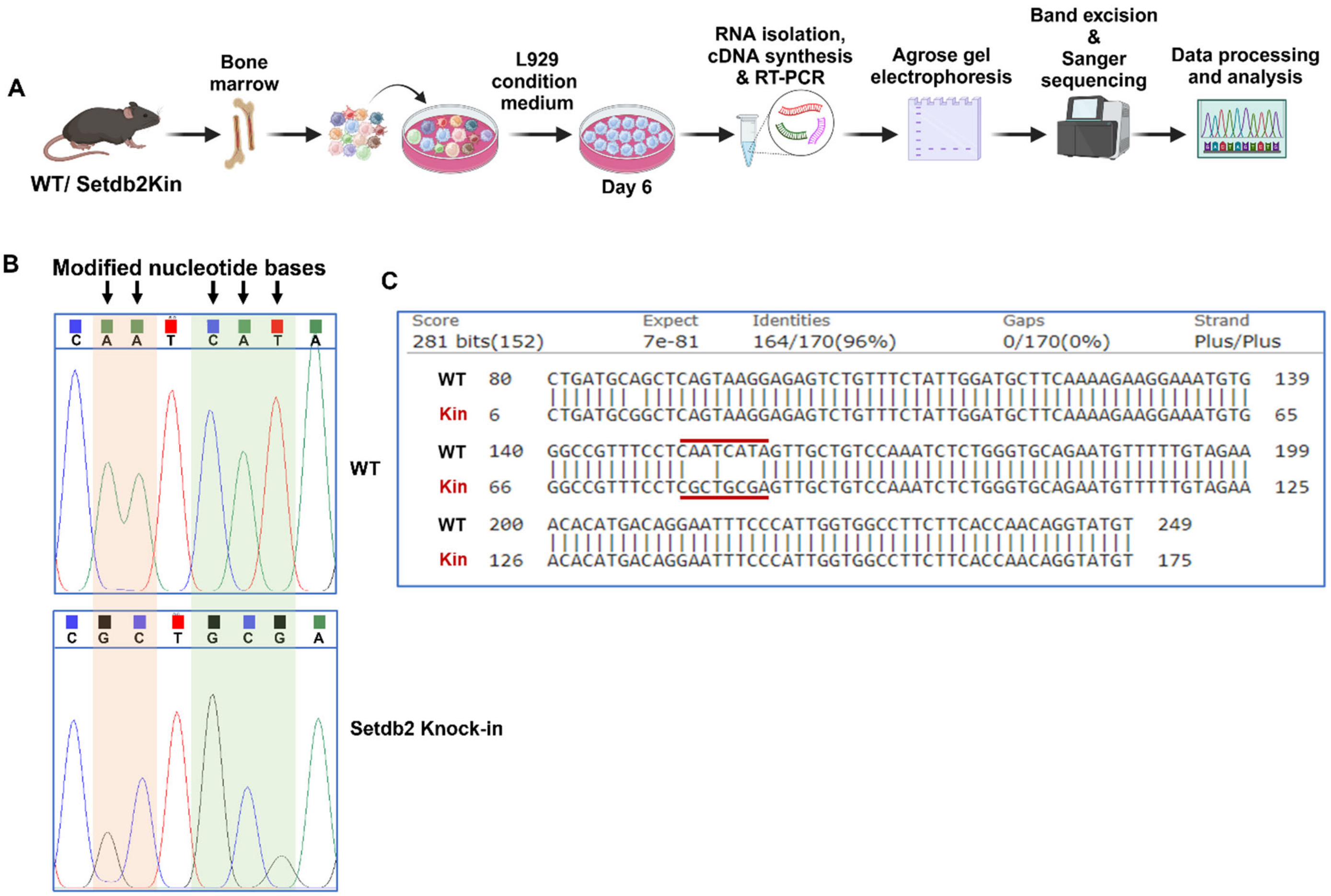

**Figure S2. Setdb2Knock-in validation through sanger sequencing in BM-macrophages (Related to Figures 4 and S6)**

(A) Work flow of Setdb2Kin confirmation. Bone marrow cells were isolated from WT and Setdb2Kin mice and differentiated with L929 condition medium for 6 days. Total RNA was isolated followed by cDNA synthesis. RT-PCR was performed with sequence specific primers (Table-S1) and products were visualized through agarose gel electrophoresis. Bands of interest were excised from the gel followed by purification and sanger DNA sequencing.

(B) Sequence chromatograms depicting targeted nucleotide bases in WT (top), and Setdb2Kin (bottom).

(C) Blast search display showing targeted nucleotide sequences in WT and Setdb2Kin mice.

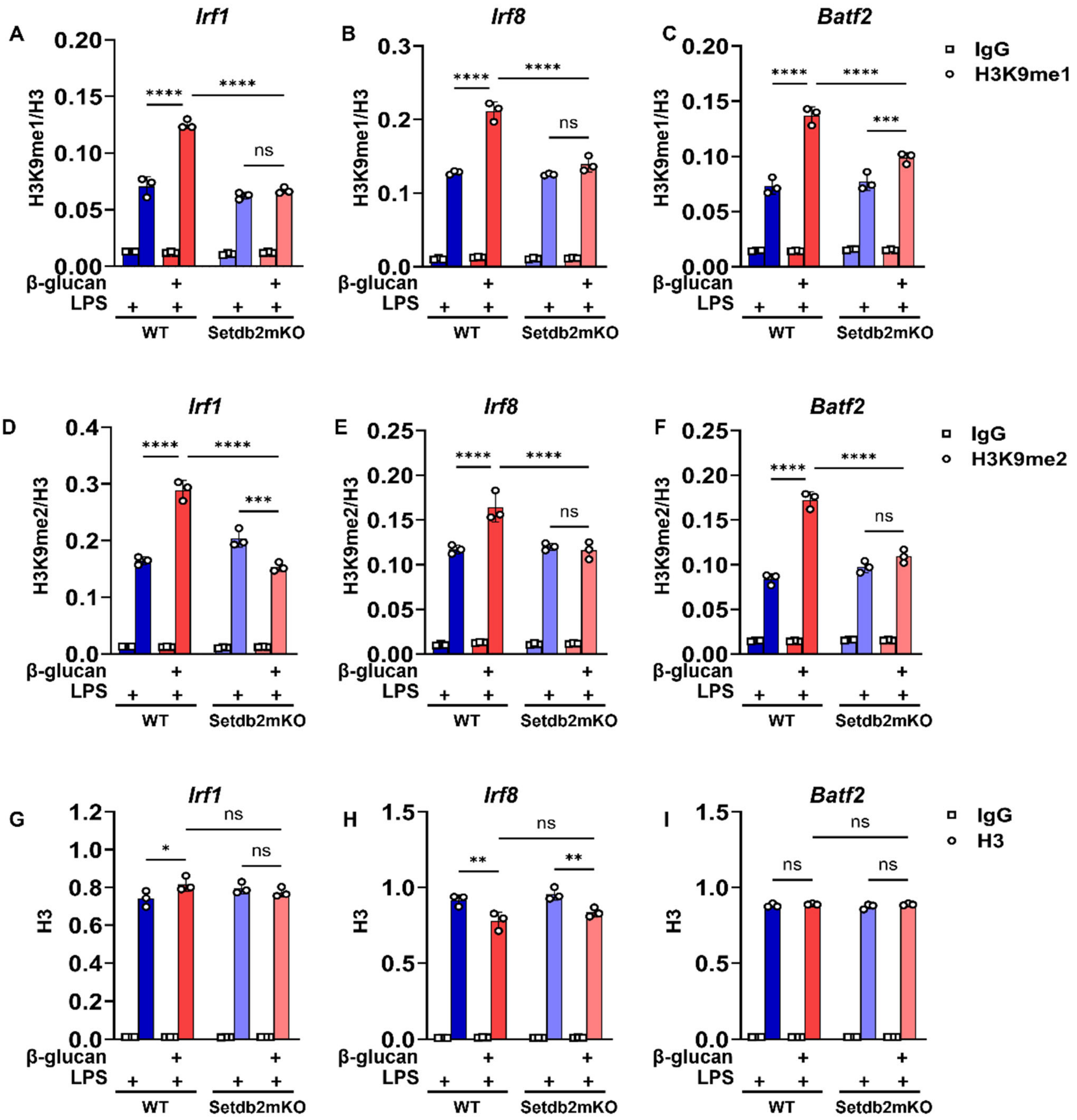

**Figure S3. H3K9me levels in β-glucan trained WT and Setdb2mKO BM-macrophages (Related to Figure 5)**
Bone marrow cells were treated as detailed in the legend of Figure 5 followed by ChIP-qPCR analysis.

(A-C) H3K9me1 (D-F) H3K9me2 and (G-I) H3 levels at *Irf1*, *Irf8*, and *Batf2* promoters in LPS alone or β-glucan followed by LPS restimulated BM-macrophages from WT (n = 3) and Setdb2mKO mice (n = 3).

Data are represented as mean ± SD. 'n' represents biological replicates of each strain and treatment type. p-value was calculated using two-way ANOVA with Tukey post-hoc test for multiple comparisons. $*p < 0.05$, $**p < 0.01$, and $***p < 0.001$, ns, not significant change.

A

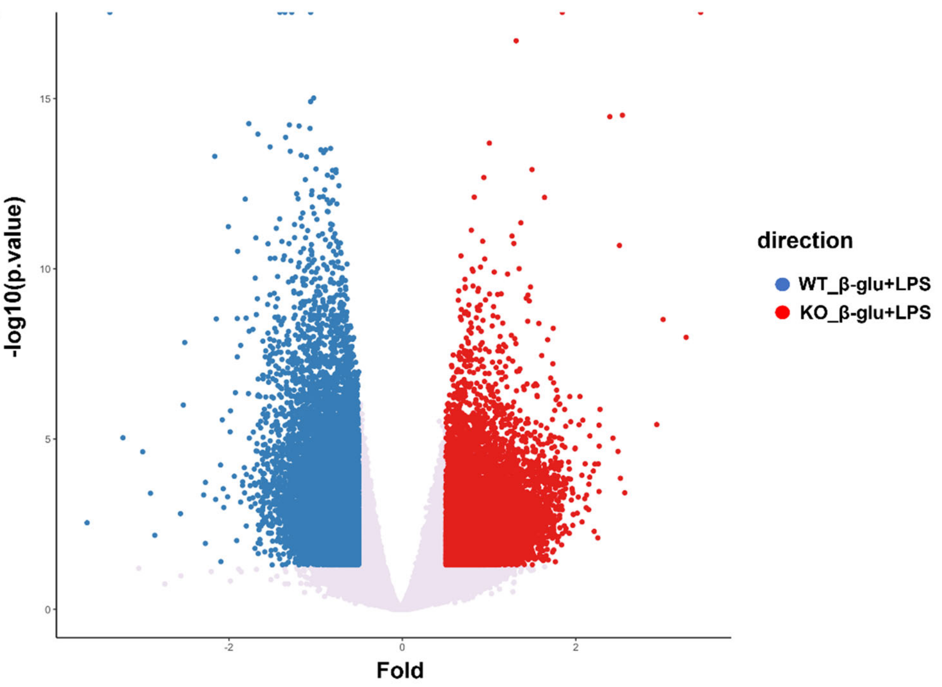
15
10
5
0
-log10(p.value)
-2
0
2
Fold
direction
WT_β-glu+LPS
KO_β-glu+LPS

B

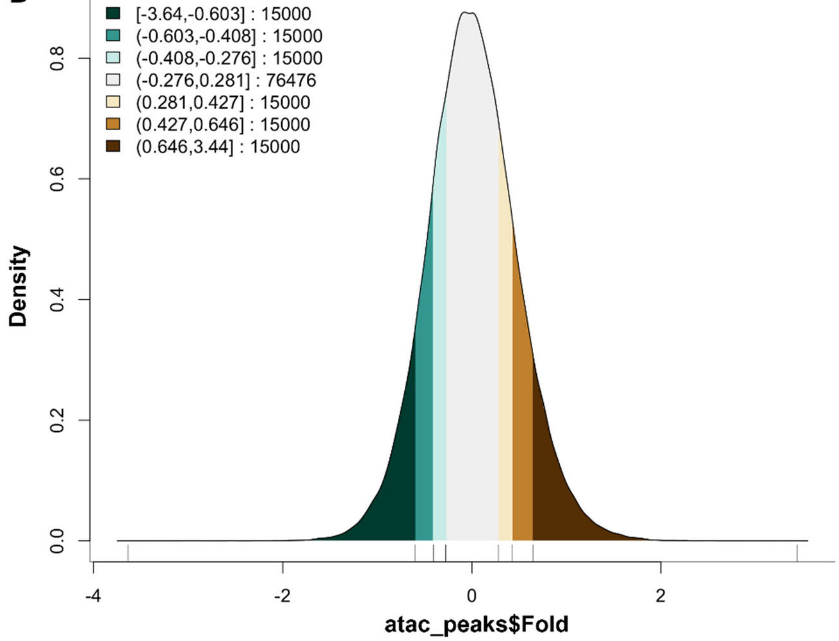
[-3.64,-0.603] : 15000
(-0.603,-0.408] : 15000
(-0.408,-0.276] : 15000
(-0.276,0.281] : 76476
(0.281,0.427] : 15000
(0.427,0.646] : 15000
(0.646,3.44] : 15000
0.8
0.6
0.4
0.2
0.0
Density
-4
-2
0
2
atac_peaks$Fold

**C**

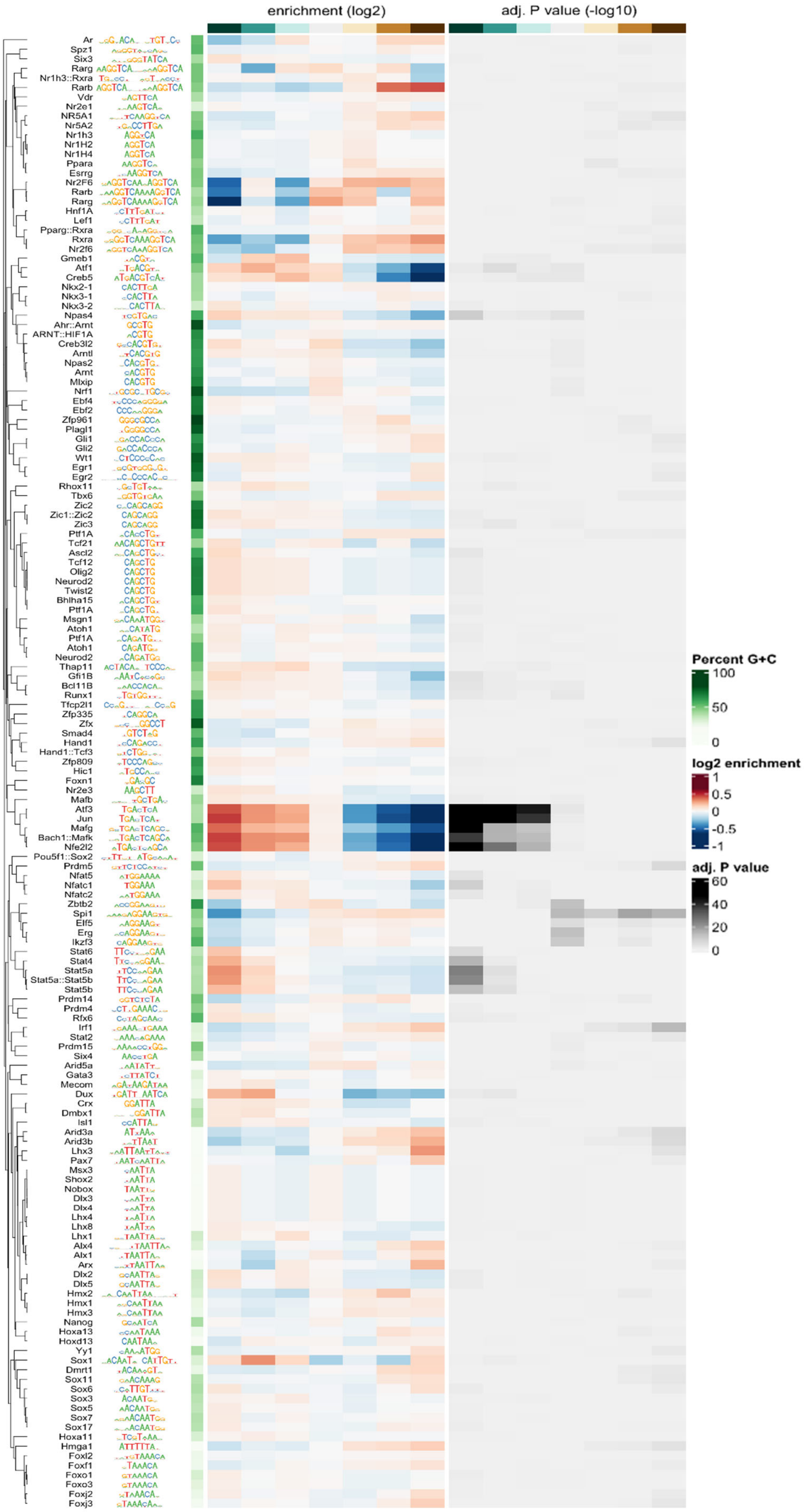
enrichment (log2)
adj. P value (-log10)
Ar
Spz1
Six3
Rarg
Nr1h3::Rxra
Rarb
Vdr
Nr2e1
NR5A1
NR5A2
Nr1h3
Nr1H2
Nr1H4
Ppara
Esrrg
Nr2F6
Rarb
Rarg
Hnf1A
Lef1
Pparg::Rxra
Rxra
Nr2f6
Gmeb1
Atf1
Creb5
Nkx2-1
Nkx3-1
Nkx3-2
Npas4
Ahr::Arnt
ARNT::HIF1A
Creb3l2
Arntl
Npas2
Arnt
Mlxip
Nrf1
Ebf4
Ebf2
Zfp961
Plagl1
Gli1
Gli2
Wt1
Egr1
Egr2
Rhox11
Tbx6
Zic2
Zic1::Zic2
Zic3
Ptf1A
Tcf21
Ascl2
Tcf12
Olig2
Neurod2
Twist2
Bhlha15
Ptf1A
Msgn1
Atoh1
Ptf1A
Atoh1
Neurod2
Thap11
Gfi1B
Bcl11B
Runx1
Tfcp2l1
Zfp335
Zfx
Smad4
Hand1
Hand1::Tcf3
Zfp809
Hic1
Foxn1
Nr2e3
Mafb
Atf3
Jun
Mafg
Bach1::Mafk
Nfe2l2
Pou5f1::Sox2
Prdm5
Nfat5
Nfatc1
Nfatc2
Zbtb2
Spi1
Elf5
Erg
Ikzf3
Stat6
Stat4
Stat5a
Stat5a::Stat5b
Stat5b
Prdm14
Prdm4
Rfx6
Irf1
Stat2
Prdm15
Six4
Arid5a
Gata3
Mecom
Dux
Crx
Dmbx1
Isl1
Arid3a
Arid3b
Lhx3
Pax7
Msx3
Shox2
Nobox
Dlx3
Dlx4
Lhx4
Lhx8
Lhx1
Alx4
Alx1
Arx
Dlx2
Dlx5
Hmx2
Hmx1
Hmx3
Nanog
Hoxa13
Hoxd13
Yy1
Sox1
Dmrt1
Sox11
Sox6
Sox3
Sox5
Sox7
Sox17
Hoxa11
Hmga1
Foxl2
Foxf1
Foxo1
Foxo3
Foxj2
Foxj3
Percent G+C
100
50
0
log2 enrichment
1
0.5
0
-0.5
-1
adj. P value
60
40
20
0

**Figure S4. ATACseq analysis in β-glucan trained WT and Setdb2mKO BM-macrophages (Related to Figure 5)**
Bone marrow cells were treated with 5 µg/mL β-glucan (training stimulus) or culture medium for 24 h, cultured for 5 days, and restimulated with 100ng/mL LPS (secondary challenge) or culture medium for 6 h on day 6, followed by ATACseq profiling and analysis.

(A) Volcano plot representing differential peaks in samples from BM-macrophages treated with β-glucan followed by LPS stimulation from WT (n = 3) and Setdb2mKO mice (n = 3). (B, C) Top enriched JASPAR motifs of differential peaks stratified by fold-change using MonaLisa (MOtif aNAlysis with Lisa) plot.

For ATAC-seq data, differential peak accessibility was analyzed using DiffBind with DESeq2, employing default parameters. Data are represented as mean ± SD. 'n' represents biological replicates of each strain and treatment type. p-value was calculated using two-way ANOVA with Tukey post-hoc test for multiple comparisons. ∗p < 0.05, ∗∗p < 0.01, and ∗∗∗p < 0.001, ns, not significant change.

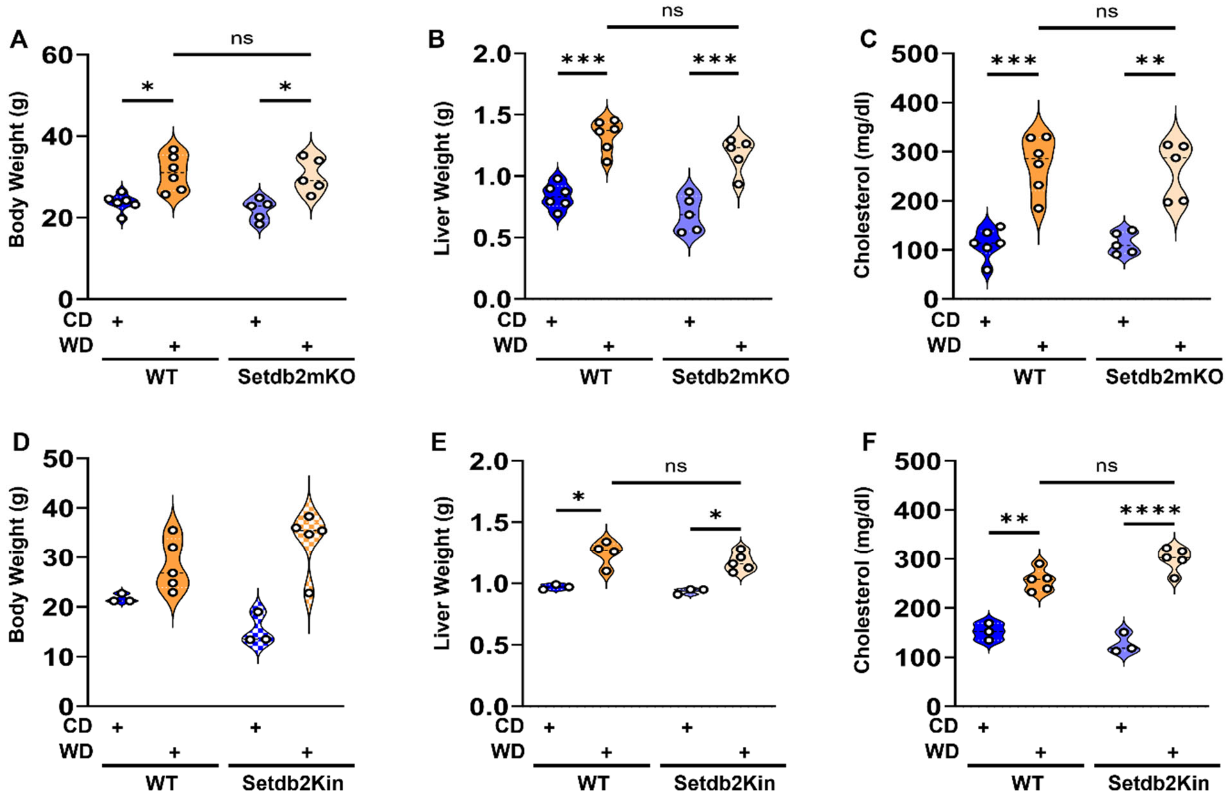

**Figure S5. Whole body assessment of control vs 'Western-diet' fed WT, Setdb2mKO and Setdb2Kin mice (Related to Figure 6 and S6)**
Mice were fed with western-diet or control-diet (training) for 20 weeks followed by estimation of body weight, liver weight and serum total cholesterol levels.

(A) Whole body weight (B) liver weight (C) and serum total cholesterol levels in control diet and western diet fed WT vs. Setdb2mKO mice. Control diet WT (n = 6), western diet WT (n = 6), control diet Setdb2mKO (n = 5), and western diet Setdb2mKO mice (n = 5).

(D) Whole body weight (E) liver weight (F) and serum total cholesterol levels in control diet and western diet fed WT vs. Setdb2Kin mice. Control diet WT (n=3), Control diet Setdb2Kin (n=3), Western diet WT (n=5), and Western diet Setdb2Kin mice (n=5). Data are represented as mean ± SD. 'n' represents biological replicates of each strain and treatment type. p-value was calculated using two-way ANOVA with Tukey post-hoc test for multiple comparisons. ∗p < 0.05, ∗∗p < 0.01, and ∗∗∗p < 0.001, ns, not significant change.

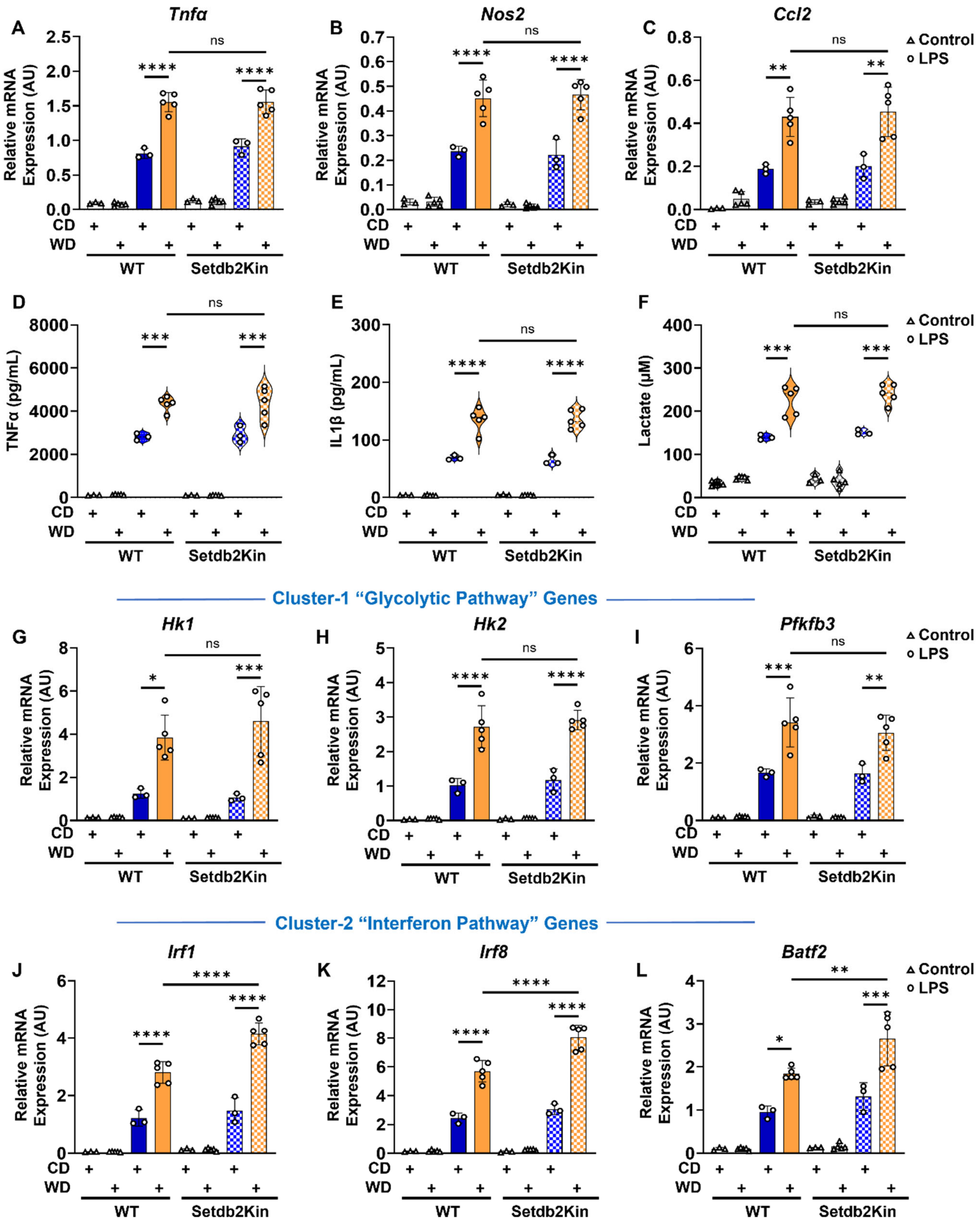
A
Tnfα
B
Nos2
C
Ccl2
Control
LPS
Relative mRNA Expression (AU)
CD
WD
WT
Setdb2Kin
ns
D
E
F
TNFα (pg/mL)
IL1β (pg/mL)
Lactate (μM)
Cluster-1 "Glycolytic Pathway" Genes
G
Hk1
H
Hk2
I
Pfkfb3
Cluster-2 "Interferon Pathway" Genes
J
Irf1
K
Irf8
L
Batf2

**Figure S6. ‘Western-diet triggered innate immune memory in BM-Monocytes from Setdb2Kin mice (Related to Figure 6 and S5)**

CD11b+ BM-monocytes were isolated from control-diet or western-diet fed WT or Setdb2Kin mice followed by stimulation with 100ng/mL LPS or culture medium in vitro for 6 h.

(A) mRNA expression of *Tnfα* (B) *Nos2*, and (C) *Ccl2*, (D) Pro-inflammatory cytokine levels of TNFα, and (E) IL-1β, (F) Extracellular lactate levels, (G) mRNA expression of *Hk1* (H) *Hk2*, (I) *Pfkfb3* (J) *Irf1* (K) *Irf8*, and (L) *Batf2* in control diet, western diet, control diet followed by *in vitro* LPS restimulation, and western diet followed by *in vitro* LPS restimulated BM-monocytes isolated from WT and Setdb2Kin mice.

Control diet WT (n=3), Control diet Setdb2Kin (n=3), Western diet WT (n=5), and Western diet Setdb2Kin mice (n=5), from one of the three independent experimental group. Data are represented as mean ± SD. ‘n’ represents biological replicates of each strain and treatment type. p-value was calculated using two-way ANOVA with Tukey post-hoc test for multiple comparisons. $*p < 0.05$, $**p < 0.01$, and $***p < 0.001$, ns, not significant change.

### Table S1

| Primer sequence for qPCR | |
|---|---|
| **Gene** | **Sequence 5'-3'** |
| *Tnfα* Forward: | CCTCACACTCAGATCATCTTCT |
| *Tnfα* Reverse: | GCTACGACGTGGGCTACAG |
| *Nos2* Forward: | CCAAGCCCTCACCTACTTCC |
| *Nos2* Reverse: | CTCTGAGGGCTGACACAAGG |
| *Ccl2* Forward: | CCCGTAAATCTGAAGCTAA |
| *Ccl2* Reverse: | CACACTGGTCACTCCTACAGAA |
| *Setdb2* Forward: | AACAAATCAAGTGCGGTTCC |
| *Setdb2* Reverse: | TTCAAGGACAGTGGGGTTTC |
| *Hk1* Forward: | AGGGCGCATTACTCCAGAG |
| *Hk1* Reverse: | CCCTGTGGGTGTCTTGTGTG |
| *Hk2* Forward: | TGATCGCCTGCTTATTCACGG |
| *Hk2* Reverse: | AACCGCCTAGAAATCTCCAGA |
| *Pfkfb3* Forward: | CCCAGAGCCGGGTACAGAA |
| *Pfkfb3* Reverse: | GGGGAGTTGGTCAGCTTCG |
| *Irf1* Forward: | ATGCCAATCACTCGAATGCG |
| *Irf1* Reverse: | TTGTATCGGCCTGTGTGAATG |
| *Irf8* Forward: | CGGGGCTGATCTGGGAAAAT |
| *Irf8* Reverse: | CACAGCGTAACCTCGTCTTC |
| *Batf2* Forward: | AGCGAGCTGCTGACTGAGA |
| *Batf2* Reverse: | CGCCTTACTGGTGTGCTTCT |
| *L32* Forward: | ACATTTGCCCTGAATGTGGT |
| *L32* Reverse: | ATCCTCTTGCCCTGATCCTT |
| **Primer sequence for ChIP-qPCR** | |
| **Gene** | **Sequence 5'-3'** |
| *Irf1* Forward: | TGCCTTGTACTTCCCCTTCG |
| *Irf1* Reverse: | GCCGCGAAGAAATCTAAACAC |
| *Irf8* Forward: | GCAAAAGTGATTTCTCGGAAAGA |
| *Irf8* Reverse: | GCTTTTATAGATGGGGCGGG |
| *Batf2* Forward: | CAGCCCATGGTTTTAGGAGAAA |
| *Batf2* Reverse: | AGAGGTTCAGTCTGGTCCTG |
| **Primer sequence for 3C-qPCR** | |
| **Gene** | **Sequence 5'-3'** |
| *Gapdh* Taqman probe | TATCCCTCCTCGGAACCTGAGAGC |
| *Gapdh* Forward: | GCAGCCTGGAAACCTGATAA |
| *Gapdh* Reverse: | GGGCTACAGTGGGTGAAAG |
| *Tnf* Taqman probe | ATTCCCAGGGCTGAGTTCATTCCC |
| Constant (*Tnf*) | GCTTGAGAGTTGGGAAGTGT |
| Test (*Tnf*) N1 | CGGAGCCCTCCAGACAT |
| Test (*Tnf*) E1 | CTGTAGCTAAATAGGACCTTGAACA |
| Test (*Tnf*) E2 | GGGTTGTCTGTTCTTGGGCA |
| Test (*Tnf*) N2 | GAGGCCATAAAGGAGGGATT |
| *Ccl2* Taqman probe | CTGTGAAGGGTTCCAACCACTCC |
| Constant (*Ccl2*) | AAACTCACCTAGTGCTTACTCTG |
| Test (*Ccl2*) N1 | CACTCTTTTTATCAAGAGTCTGCTG |
| Test (*Ccl2*) E1 | GCTGCCTCCTGTGCTAA |
| Test (*Ccl2*) N2 | AAGACCACCTATCTAGTTACAAGTG |
| *Hk2* Taqman probe | CACGGAACACACGTCCCAACTCT |
| Constant (*Hk2*) | AGCCAATCAGCGCCTAGAG |
| Test (*Hk2*) N1 | TGCTGACGATGGCGCTAA |
| Test (*Hk2*) E1 | TCAGGTGGGAAATTCCTTCCTT |
| Test (*Hk2*) N2 | GAGTTCTAGCCTAGAGCAGCA |
| **Primer sequence for sanger sequencing** | |
| **Gene** | **Sequence 5'-3'** |
| *Setdb2kin* Forward: | TGCCAAGTGAGAGGACAAAG |
| *Setdb2kin* Reverse: | ACATACCTGTTGGTGAAGAAGG |
